# A Single Site Mutation Tunes Fluorescence and Chromophorylation of an Orange Fluorescent Cyanobacteriochrome

**DOI:** 10.1101/2023.05.11.540396

**Authors:** Makena K. Janis, Wenping Zou, Melissa L. Zastrow

**Author notes:** These authors contributed equally to this work (Makena K. Janis, Wenping Zou).

## Abstract

Cyanobacteriochrome (CBCR) GAF domains bind bilin cofactors to confer sensory wavelengths important for various cyanobacterial photosensory processes. Many isolated GAF domains autocatalytically bind bilins, becoming fluorescent. The third GAF domain of CBCR Slr1393 from *Synechocystis* sp. PCC6803 binds phycocyanobilin (PCB) natively, yielding red/green photoswitching properties but also binds phycoerythrobilin (PEB). GAF3-PCB has low quantum yields but non-photoswitching GAF3-PEB is brighter, making it a promising platform for new genetically encoded fluorescent tools. GAF3, however, shows low PEB binding efficiency (chromophorylation) at ∼3% compared to total protein expressed in *E. coli*. Here we explored site-directed mutagenesis and plasmid-based methods to improve GAF3-PEB binding and demonstrate its utility as a fluorescent marker in live cells. We found that a single mutation improved chromophorylation while tuning the emission over ∼30 nm, likely by shifting autoisomerization of PEB to phycourobilin (PUB). Plasmid modifications also improved chromophorylation and moving from a dual to single plasmid system facilitated exploration of a range of mutants via site saturation mutagenesis and sequence truncation. Collectively, the PEB/PUB chromophorylation was raised by ∼7-fold. Moreover, we show that protein-chromophore interactions can tune autoisomerization of PEB to PUB in a GAF domain, which will facilitate future engineering of similar GAF domain-derived fluorescent proteins.

## Introduction

Cyanobacteriochromes (CBCRs) are a group of phytochrome photoreceptors that sense and respond to light to control a range of physiological and metabolic processes in cyanobacteria.^1^ Unlike canonical phytochromes, cyanobacteriochromes require only the chromophore-binding GAF (cGMP-specific phosphodiesterase, adenylyl cyclase, and FhlA protein) domain for photoconversion.^2–4^ All phytochromes use linear tetrapyrrole chromophores (bilins) as the photosensory elements.^5^ In CBCRs, the bilin cofactor is usually covalently attached to a conserved cysteine residue within the GAF domain via a thioether bond to the C3 side chain of the bilin A ring.^3^ Four different bilin chromophores have been found natively in CBCR GAF domains, phycocyanobilin (PCB), phycochromobilin (PΦB), phycoviolobilin (PVB), and likely biliverdin (BV).^6–10^ These bilin cofactors can photoisomerize between 15Z and 15E configurations at the double bond to the D-ring.^4^ Gene *slr1393* from cyanobacterium *Synechocystis* sp. PCC6803 includes three GAF domains, and the GAF3 domain alone auto-catalytically binds PCB as the native cofactor.^11–13^

CBCRs are promising platforms for engineering new fluorescent proteins. Compared to other phytochromes, they require only a single small domain for autocatalytic bilin attachment. The photoconverting nature of many bilin-binding GAF domains, however, is associated with low quantum yields for both photostates.^14^ High quantum yields can be achieved by using a non-photoconverting bilin cofactor like phycoerythrobilin (PEB).^15^ Slr1393-GAF3 (hereafter referred to as GAF3) assembled with the PCB chromophore shows reversible photoconversion between green (λ_max/abs_ = 650 nm) and red (λ_max/abs_ = 539 nm) states with quantum yields of 6% and 3%, respectively.^11,13^ GAF3 can also bind PEB.^15^ The lack of a Δ15, 16 double bond between pyrrole ring C and D in PEB removes the ability to photoswitch resulting in higher quantum yield (31%) with orange fluorescence (Table 1, Scheme S1).^15^

**Table 1.** Spectroscopic properties of PEB/PUB-bound Slr1393-GAF3 and GAF3 mutants.

| Protein | Absorption |  | Fluorescence |  |
| --- | --- | --- | --- | --- |
| | $\lambda_{\text{max}}$ (nm)<sup>[a]</sup> | $\epsilon$ (mM<sup>-1</sup>cm<sup>-1</sup>)<sup>[b]</sup> | $\lambda_{\text{max}}$ (nm)<sup>[c]</sup> | $\Phi$ |
| GAF3<sup>[d]</sup> | 530 | 42.8 | 575 | 0.31 |
| GAF3<sup>[e]</sup> | 507 (w), 530, 567 (w) | 200 ± 30, 42.8 ± 2.8, 36 ± 4 | 560/580 | n.d. |
| GAFW496Y<sup>[e]</sup> | 505, 532, 560 | 200 ± 20, 48 ± 4, 46 ± 4 | 578/580 | 0.43 ± 0.07 |
| GAF3W96H<sup>[e]</sup> | 503, 522, 567 (w) | 150 ± 20, 44 ± 6, 28 ± 4 | 552/582 | 0.56 ± 0.09 |
| trGAF3<sup>[e]</sup> | 506 (w), 530, 565 (w) | 151 ± 19, 37 ± 1, 29 ± 1 | 559/580 | 0.51 ± 0.02 |
| trGAFW496Y<sup>[e]</sup> | 505 (w), 533, 562 | 180 ± 9, 46 ± 3, 44 ± 2 | 580/582 | 0.43 ± 0.03 |
| trGAF3W496H<sup>[e]</sup> | 505, 522, 567 (w) | 205 ± 6, 46 ± 1, 26 ± 2 | 551/579 | 0.46 ± 0.03 |
| trGAF3W496V<sup>[e]</sup> | 502, 532 | 214 ± 6, 53 ± 1 | 558/584 | 0.43 ± 0.05 |
| trGAF3W496G<sup>[e]</sup> | 506, 532 | 237 ± 8, 53 ± 2 | 560/580 | 0.38 ± 0.01 |

[a] “w” denotes weak or shoulder peaks. [b] Extinction coefficients for wavelengths around 500-507 were determined using the free PUB extinction coefficient as the standard and those from ~530-567 nm were determined using the free PEB extinction coefficient as the standard (further details are provided in the Experimental Section). [c] The first $\lambda_{\text{max}}$ value is obtained from excitation at 490 nm and the latter from excitation at 520-550 nm. Presumably these values represent the PUB-bound and PEB-bound fractions of the protein, respectively. [d] Data from reference <sup>15</sup>. [e] Data collected in this work.

Bilin-binding GAF domains can be co-expressed in *E. coli* with bilin synthesis enzymes that produce PCB, PEB, or other bilins. Biosynthesis of these bilins begins with degradation of heme by heme oxygenase to yield BV (Scheme S1).^16^ In cyanobacteria, BV is further metabolized to yield PCB, PΦB, or PEB by ferredoxin-dependent bilin reductases. The higher quantum yield and lack of photoswitching for GAF3-PEB make it a promising bilin-binding fluorescence protein for designing new biosensors. Although other PEB-binding phycobiliproteins like phycoerythrin, phycocyanin, and allophycocyanin have been elucidated, the required extra lyase for chromophore assembly limits their application.^17,18^ Autocatalytic PEB assembly in the GAF3 domain together with the smaller size of GAF3 (∼18 kDa) outperforms other phycobiliproteins.

To fully harness GAF3-PEB as a fluorescent biomarker, however, the PEB binding efficiency must be improved. Native Slr1393-GAF3 binds PCB with a chromophorylation ratio of 61% when co-expressed with PCB synthesis enzymes heterologously in *E. coli* (Table S1). In contrast, PEB has a far lower binding efficiency of ∼3% to GAF3, making its application for live cell imaging challenging. Upon binding, PEB also partially autoisomerizes to the less conjugated phycourobilin (PUB) cofactor (Scheme S1), but the effects of the protein structure on this activity are unknown and vary among different GAF domains.^15^ The effects of protein-chromophore interactions on photoswitching have been explored in various PCB, PΦB, PVB, and BV-binding GAFs and phytochromes, but such effects on PEB-binding GAF protein spectroscopic properties have not been well-studied.^19–24^

Here we took two approaches to improving the chromophorylation and tuning the spectroscopic properties of GAF3-PEB. We first identified two site-directed mutants that each led to ∼2-fold increase in chromophorylation and varied maximum emission wavelength by ∼30 nm. Using the red-shifted mutant, GAF3_W496Y_, we explored several plasmid-based approaches to improving PEB chromophorylation by modulating the relative expression levels of GAF3 and the PEB synthesis enzymes. After achieving nearly 4-fold higher PEB binding, we used a single plasmid approach combining all PEB synthesis enzymes and GAF3 in one expression plasmid. The single plasmid facilitated protein truncation and site-directed mutagenesis, further improving chromophorylation up to 7-fold and modulating the spectroscopic properties of GAF3, likely by modifying the extent of autoisomerization of GAF3-PEB to GAF3-PUB. With these collective efforts we tuned PEB chromophorylation from ∼3-19% and achieved a range of fluorescence emission maxima from ∼550 to 580 nm. We also show that truncated GAF3 proteins can be readily used to image live *E. coli* cells with fluorescence microscopy. Based on these results, truncated GAF3 and mutants represent a series of orange fluorescent proteins suitable for use as biomarkers and in future biosensing scaffolds.

## Results and Discussion

### Cloning and Protein Expression of GAF3 and GAF3 Mutants with Bilin Chromophores

The plasmids encoding GAF3 and PCB or PEB synthesis enzymes were prepared similarly to those previously published (Supporting Information).^11,25,26^ Briefly, the pET28a-GAF3 plasmid encodes the gene for His_6_-tagged Slr1393-GAF3 protein. GAF3 mutants were prepared by site-directed mutagenesis of pET28a-GAF3 (primers listed in Table S2). The *ho1* and *pcya* or *ho1* and *pebs* genes were encoded in the plasmid pACYCDuet (Table S3). For protein expression, pET28a-GAF3 and pACYCDuet-HO1-PcyA or pACYCDuet-HO1-PebS were co-transformed for GAF3-PCB or GAF3-PEB production in *E. coli*. After co-expression and protein purification (Supporting Information), GAF3-PEB protein was used for spectral analyses. Each GAF3 mutant was co-expressed with the enzymes for PEB synthesis and isolated using the same procedure. Protein purity was confirmed by denaturing gel electrophoresis (Figure S1).

### Bilin Binding in GAF3 and GAF3 mutants

The UV-visible and fluorescence spectra of GAF3-PCB and GAF3-PEB are similar to those previously reported (Figure 1, S2). The full UV-visible spectrum of GAF3-PEB, however, revealed low binding of PEB to GAF3 (i.e. chromophorylation). By comparing the absorbance at 280 nm (ε = 34950 M^-1^ cm^-1^) with the absorption of bound PEB at 530 nm (ε = 42800 M^-1^ cm^-1^), the calculated chromophorylation is 3% (Table S1, Experimental Section). Including the percentage of PEB autoisomerized to PUB (λ_max_ = 507 nm, ε = 200000 M^-1^ cm^-1^, Table 1) raises this number by only ∼0.6%. By comparison, the PCB chromophorylation of GAF3 is 61% when both the Z and E photoisomerization states are accounted for (ε_648_ = 97000 M^-1^ cm^-1^ and ε_556_ = 47000 M^-1^ cm^-1^).^11^ We initially hypothesized that the change in structure between PCB and PEB, whereby the C15-C16 double bond in PCB becomes saturated in PEB, may negatively affect the chromophorylation by interrupting stabilizing interactions around the C and D ring in the binding site (Scheme S1). Other studies have successfully converted the selectivity of CBCRs to bind non-cognate bilins such as BV with up to 75% chromophorylation using a rational site-directed mutagenesis approach.^27^ Prior work on GAF3 shows that the side chain indole group on tryptophan 496 has a π-π stacking interaction with pyrrole ring D of PCB.^11,12^ This residue is also known as the “lid” tryptophan in related CBCR red/green photoswitchers, AnPixJg2 and NpR6012g4, although mutation in the latter does not interrupt the red/green photocycle.^28,29^ X-ray crystal structures of both photostates in AnPixJg2 and Slr1393-GAF3 show that this Trp undergoes a significant shift in position during photoisomerization of PCB.^28,30^ The C15-C16 double bond in PCB restricts free rotation about the bond. In PEB, however, the C15-C16 single bond can undergo free rotation, and the presence of Trp may lead to steric clashing. Since covalent bilin attachment occurs via initial reversible non-covalent binding of the bilin within the GAF pocket followed by reaction with Cys to form the thioether bond,^31^ Trp may hinder the first non-covalent binding step of PEB. After analyzing structure alignments between PCB-bound Slr1393-GAF3 (PDB 5DFX), a PEB-bound GAF1 domain from All2699 of *Synechocystis* sp. PCC 7120 (PDB 6OZB), and PEB-containing phycobilins, like B-phycoerythrin from *Porphyridium cruentum* (PDB 3V58) and phycoerythrin from *Gloeobacter violaceus* (PDB 2VJH), we found a tyrosine side chain in each structure oriented toward the phycobilin cofactor in a similar position to GAF3 Trp496 (Figure S3A-B).^30,32,33^ We also compared sequence alignments between GAF3 and several other CBCRs and related phytochromes (Figure S4). Red/green CBCRs retain Trp in the position analogous to W496. Many related phytochromes, however, have a Tyr residue in this position in the alignment. In All2699, a Phe residue (F83) is present, although the crystal structure reveals that a different tyrosine (Y142) is positioned closer to where the GAF3-W496 residue is overlaid (Figure S3A and S4). In the phytochrome Cph1, which can also bind PEB,^14,31^ a Tyr (Y203) is present in the sequence alignment. An overlay of the crystal structures of Cph1-PCB and GAF3-PCB reveals that this Tyr lies further from the D ring in Cph1 than W496 in GAF3, but both are on flexible loop regions (Figure S3C). We also compared a CBCR GAF domain that served as a template for re-engineering bilin binding from PCB to BV to generate the near-infrared fluorescent protein miRFP670nano.^34,35^ Here, the residue in the position analogous to GAF3-W496 is valine. Examination of modeled Trp496 mutants in GAF3-PCB revealed that mutation may increase space in the cofactor-binding pocket, possibly better accommodating PEB (Figure 1A-C). We tested two mutations, Trp496 to Tyr and to His, both of which could potentially retain some π-π stacking interactions while removing steric hindrance (Figure 1A-C).

**Figure 1.**
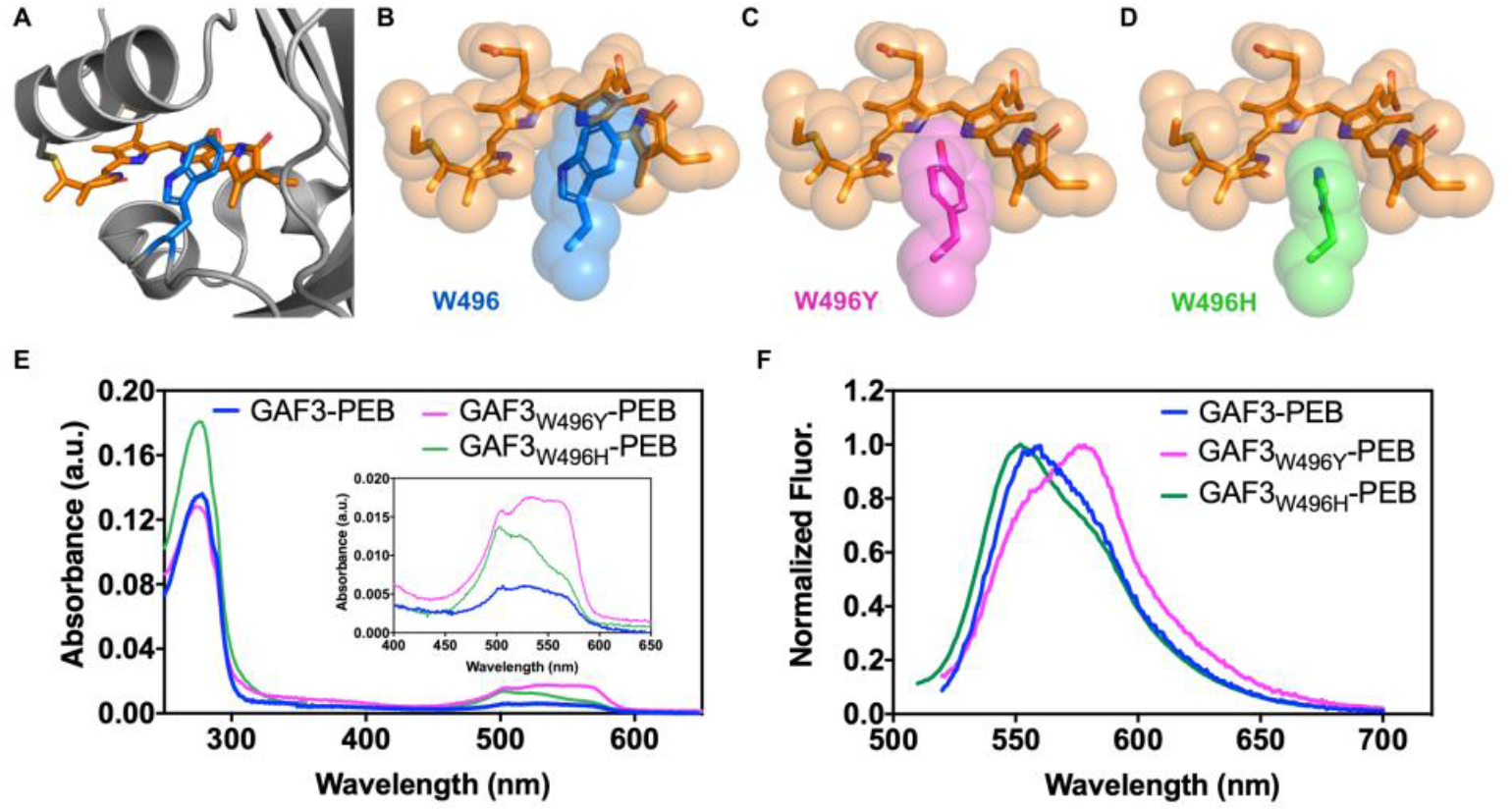
Structures and UV-visible and fluorescence spectroscopy of bilin-bound Slr1393-GAF3. A) PyMOL model of the crystal structure of GAF3-PCB in the red state (PDB 5DFX). The protein structure is gray, the PCB cofactor is orange, and the side chain of the tryptophan-496 mutated is blue. B) Space-filling model of PCB and Trp496 from part A. Space-filling PyMOL models of the bilin-binding site in C) GAF3_W496Y_ and D) GAF3_W496H_. Mutations were modeled using PyMOL mutagenesis wizard. The tyrosine side chain in GAF3_W496Y_ is colored magenta and the histidine side chain in GAF3_W496H_ is colored green. E) UV-visible absorption and F) fluorescence spectra of GAF3-PEB (blue), GAF3_W496Y_-PEB (magenta), and GAF3_W496H_-PEB (green). Fluorescence spectra were normalized to maximum emission for each protein (Table 1). Protein concentrations were ∼3–5 μM. λ_ex_ = 490 nm. Buffer = 50 mM HEPES, 100 mM NaCl, pH 7.1

The spectroscopic properties of GAF3, GAF3_W496Y_, and GAF3_W496H_ are distinct (Figure 1E-F). As previously reported, the GAF3-PEB absorption spectrum shows that some PEB isomerizes to PUB (Scheme S1), which is characterized by absorption around 500 nm.^15^ The absorbance maxima of PEB are usually ∼530-560 nm. All absorption spectra show peaks or shoulders around 503-507 nm, 522-532 nm, and 560-567 nm, but with varied relative intensities (Figure 1E, Table 1). The most prominent absorption for GAF3-PEB is a broad band centered around 530 nm. In GAF3_W496Y_-PEB, the shoulder around 560 nm is increased and that around 505 nm is decreased compared to GAF3-PEB. This change in relative absorption maxima between GAF3-PEB and GAF3_W496Y_-PEB indicates that the PEB to PUB ratio may be increased, assuming that the absorbance from ∼530-570 nm originates from PEB (Figure 1E). Conversely, the PEB to PUB ratio significantly decreases in GAF3_W496H_, judging by the ∼2-fold higher absorbance at 505 nm compared to that from ∼520-570 nm. Using absorbance maxima and extinction coefficients corresponding to PEB and PUB for the three GAF3 proteins, we quantified and compared bound PEB vs. PUB (Table 1, S1). The PEB binding efficiency of both mutants was improved by ∼2-fold to 6 ± 2% for GAF_W496Y_ and to 5 ± 1% for GAF3_W496H_ compared to 3% for GAF3. Consistent with the relative absorption maxima, the amount of PUB formed is ∼0.6% out of 3.6% total bilin bound in GAF3, but less in GAF3_W496Y_ at ∼1% PUB out of 7% total bilin, and more in GAF3_W496H_ at 11% PUB out of 16% total bilin. The fluorescence spectra are consistent with the varied PEB to PUB ratios for GAF3 and the mutants (Table 1, Figure 1F, S5). With fluorescence excitation at 490 nm, GAF3-PEB emits with a maximum at 560 nm, but as the excitation wavelength is varied up to 560 nm, the emission shifts to 580 nm, presumably due to excitation of PEB over PUB. For GAF3_W496H_, an emission peak at 552 nm is most prominent with λ_ex_ of 490 nm, and for GAF3_W496Y_, emission at 578 nm is most prominent with the same excitation. As for GAF3, increasing the excitation wavelength shifts the emission further to PEB for both mutants. Excitation spectra highlight both PUB (∼520 nm) and PEB (∼570 nm) peaks in all mutants. The relative intensities correspond with the PEB/PUB ratio increasing in GAF3_W496Y_ and decreasing in GAF3_W496H_ relative to GAF3.

These results show that site-directed mutagenesis of Trp496 is important for improving PEB binding efficiency, possibly due to reduced steric hindrance. Furthermore, mutation at this site tunes the emission maxima in an excitation wavelength-dependent manner (Figure S5). Given that variation of the fluorescence excitation wavelength yields emission spectra with peak maxima ranging from ∼550 to 580 for both GAF3 and GAF3_W496H_ but mostly at ∼580 nm only for GAF3_W496Y_, we hypothesize that position 496 plays a role in tuning the autoisomerization ratio between bound PEB (∼580 nm) and PUB (∼550 nm).

This result may be due to varied steric packing of W496, Y496, and H496 with the chromophore or variations in π-π stacking interactions with the D ring.

### Plasmid Editing for Tuning Relative Expression of GAF Proteins and Bilin Synthesis Enzymes

Next, we explored plasmid-based methods to determine whether altering relative expression levels of GAF protein and PEB synthesis enzymes could affect chromophorylation. GAF3_W496Y_ was used here because this mutant showed two-fold more chromophorylation than GAF3 and was more red-shifted than the similarly improved GAF3_W496H_ mutant. For expression of GAF3 and GAF3 mutants described thus far, the *gaf3* gene is encoded in a pET plasmid and both *ho1* and *pebs* genes for PEB synthesis are encoded in the pACYCDuet plasmid (Figure 2A).^11,15,25^ We hypothesized that the amount of PEB produced by HO1 and PebS enzymes expressed in the cells could be a limiting factor in yielding complete GAF3-PEB. Low expression of the enzymes due to challenges in expressing multiple genes from a single plasmid could result in low amounts of PEB chromophore and limit PEB-bound protein yields. Expression of multi-protein complexes or multiple enzymes can be influenced by plasmid copy number, ribosome binding site (RBS) sequence, and rare codon usage.^36–39^ Controlled co-expression of up to four enzymes in *E. coli* BL21(DE3) cells has been achieved using T7 promoters and terminators for each gene in tandem sequence.^40^ Gene position in the plasmid and plasmid copy number can also affect multi-protein expression. Here we encoded *pebs* and *ho1* genes with a T7 promoter and terminator for each gene in the high copy number (∼40) pET28a plasmid (pET28a-PebS-HO1, Figure 2A). This plasmid was cloned using a combination of overlap extension PCR and Gibson Assembly approaches (Supporting Information, Table S4, Figure S6-S7). The *pebs* gene was placed upstream to the *ho1* gene. The gene encoding GAF3_W496Y_ was cloned into the lower copy number (∼5-10) pACYCDuet plasmid using restriction enzymes to make pACYCDuet-GAF3_W496Y_ (Supporting Information, Table S4, Figure 2A). The plasmids pACYC-GAF3_W496Y_ for GAF3_W496Y_ expression and pET28a-PebS-HO1 for PEB synthesis were then co-transformed into *E. coli* BL21(DE3) cells for expression of GAF_W496Y_-PEB protein. The UV-visible spectrum of purified GAF_W496Y_-PEB revealed an improved PEB binding value of 11 ± 2% (Figure S8, Table S1). The total protein yield, however, decreased by ∼10-fold to 0.8 mg/L from ∼7.5 mg/L for GAF_W496Y_-PEB and is likely a result of the relatively low copy number of the pACYCDuet plasmid.

**Figure 2.**
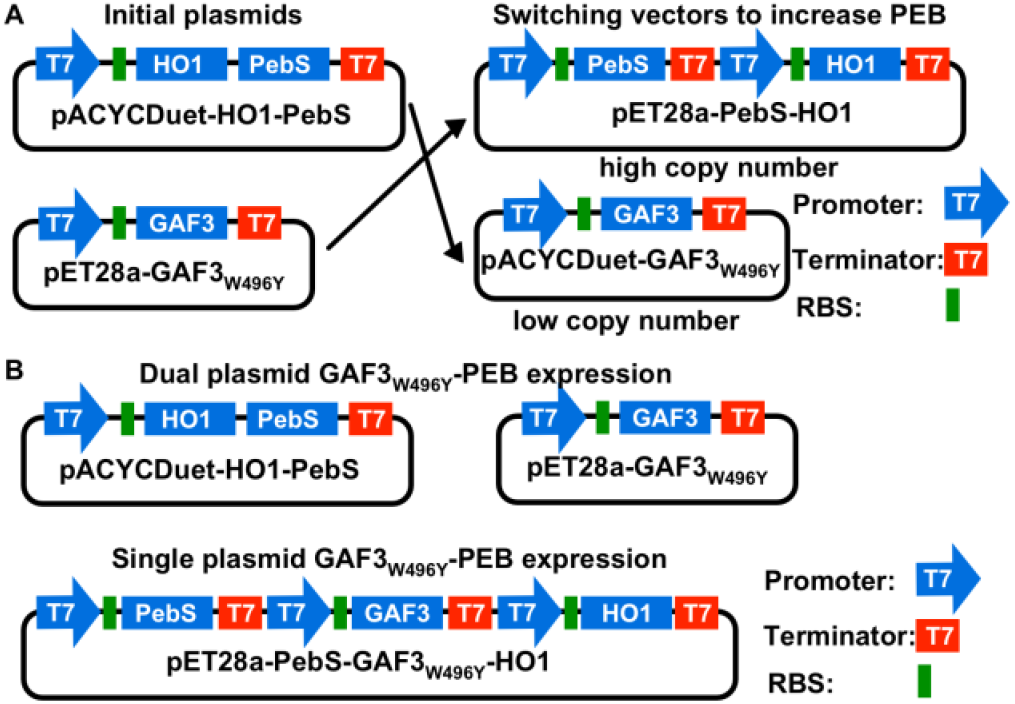
Plasmid maps for (co)expression of GAF3_W496Y_ and PEB synthesis enzymes. A) Plasmid maps of pACYCDuet-HO1-PebS and pET28a-GAF_W496Y_ used initially compared to plasmids pET28a-PebS-HO1 and pACYCDuet-GAF3_W496Y_ where varied copy numbers may adjust the relative expression levels of PEB synthesis enzymes and GAF3_W496Y_. B) Plasmid maps for dual plasmid and single plasmid expression of GAF3_W496Y_-PEB

The low yield of GAF3_W496Y_ expressed from pACYCDuet makes it challenging to isolate useful quantities of GAF3_W496Y_-PEB. We therefore encoded *ho1* and *pebs* genes back into the pACYCDuet plasmid but retained the additional T7 promoter, RBS, and T7 terminator (Supporting Information, Table S5, Figure S9A), and returned to the high copy number pET28a-GAF3_W496Y_ plasmid. GAF3_W496Y_ expressed from the pET plasmid, however, when combined with pACYCDuet-PebS-HO1 may exceed the cellular translational machinery capacity and reduce expression of PEB synthesis enzymes.^41^ We hypothesized that GAF3_W496Y_ expression could be adjusted by mutating the RBS sequence to increase the amount of PEB relative to GAF3_W496Y_, and thereby produce more PEB-bound GAF3_W496Y_ protein product. RBS variants of plasmid pMH1102 were previously investigated and pMH48, the plasmid with the second lowest translation initiation rate (TIR, used to estimate the mRNA translation rate), showed the highest chromophore binding efficiency of PCB with Cph1.^41^ We compared pMH48, pMH51, and pMH52 plasmid systems for GAF3_W496Y_-PEB production. pMH48 was chosen for the relatively low reported TIR (8164 a.u.) and pMH51 and pMH52 were selected to allow for a comparison across a range of TIR values (99029 a.u. for pET28a, 30658 a.u. for pMH51, and 48957 a.u. for pMH52). Characterization of isolated protein from each plasmid shows that the pMH51 co-expression system yielded the highest PEB binding efficiency, 9 ± 2%. PEB loading for GAF3_W496Y_ co-expressed using pMH52 was low (2.3%, Table S1, Figure S9B). The GAF3_W496Y_ yield from pMH48 was too low for isolation and measurement of PEB binding (data not shown).

Given the modest chromophorylation improvements from mutagenesis and plasmid modification approaches described thus far, further improvement to GAF3 to increase PEB binding likely requires screening more mutants, which would be facilitated by a single plasmid expression system. Furthermore, improvements to plasmid stability by using a single plasmid over a dual plasmid system for protein expression of PCB- and PEB-binding proteins that require a lyase have previously helped raise chromophorylation efficiency.^42^ Therefore, we combined genes *gaf3*_*W496Y*_, *pebs*, and *ho1* all into pET28a along with T7 promoters and terminators for each gene to generate plasmid pET28a-PebS-GAF3_W496Y_-HO1 (Figure 2B, Supporting Information, Table S5). Chromophorylation of GAF_W496Y_-PEB expressed from this single plasmid was ∼7% (Figure S10, Table S1), which is similar to that of GAF_W496Y_-PEB isolated from the dual plasmid expression system (pET28a-GAF3_W496Y_ and pET28a-PebS-HO1, 6%) and remains ∼2-fold higher than wildtype GAF3-PEB isolated from the dual plasmid system.

### Truncation and Site-Saturation Mutagenesis of GAF3

With a single plasmid system to facilitate GAF3-PEB expression in *E. coli*, we further explored sequence-based approaches to improve chromophorylation. First, we analyzed the overall structure and considered whether changes to the N- or C-termini could improve the chromophorylation by reducing protein-protein interactions that could lead to aggregation. By comparing the GAF3 crystal structure with that of another CBCR that had been engineered as a fluorescent protein (miRFP670nano), a portion of the sequence at the C-terminus was identified that was not present in miRFP670nano or the template for miRFP670nano (Figure 3A-B).^34,35^ This sequence fragment extends the C-terminal α-helix by about one heptad and given the location might promote unwanted interactions between GAF domains and possibly aggregation. We have found that homology models of various GAF domains with this heptad are often predicted as dimers via helix-helix interactions (not shown). With pET28a-PebS-GAF3_W496Y_-HO1 as the template, PCR was used to delete codons for the last eight residues and add a codon for serine (Table S6). After expression, truncated GAF3_W496Y_ (hereafter referred to as trGAF3_W496Y_) was isolated and a slightly improved (albeit not statistically significant) chromophorylation ratio was measured (∼9%, Figure 3C, Table S1). The UV-visible and fluorescence spectra of trGAF3_W496Y_-PEB are similar to those of GAF3_W496Y_-PEB (Figure 3C-D, Table 1).

**Figure 3.**
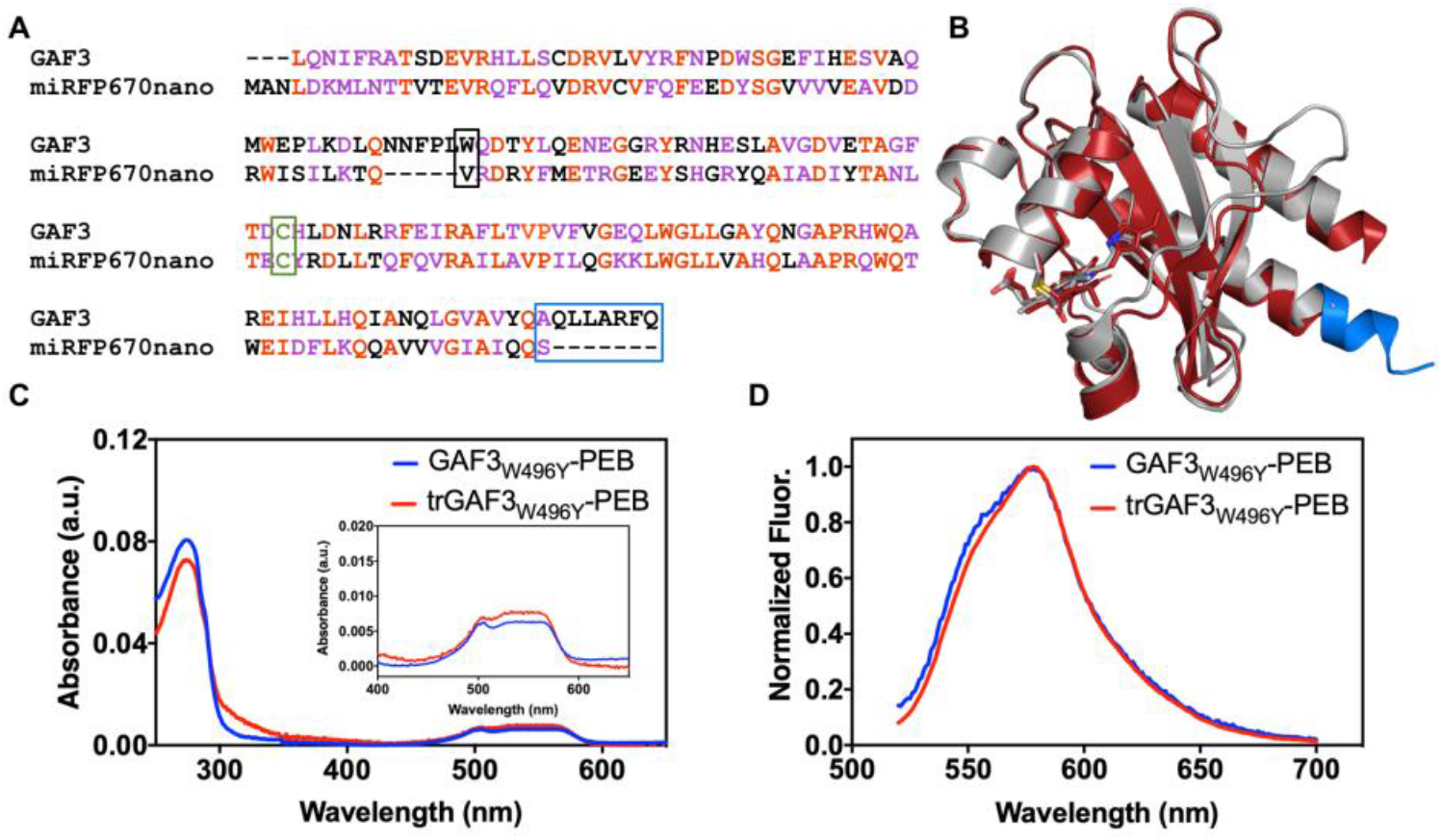
Truncation of GAF3_W496Y_. A) Sequence alignment of GAF3 (amino acids #441-596 from Slr1393) and miRFP670 (#17-163). Aligned residues are red and similar residues are purple. The conserved cysteine (C648 in GAF3) that binds to the bilin in both GAFs is green and outlined. The W496 position and corresponding residue in miRFP670nano (valine) are outlined in black. The C-terminal region truncated in GAF3 is outlined in blue. A serine codon was added after the C-terminal Gln codon. B) Overlay of PyMOL representations of the crystal structure of Slr1393 GAF3-PCB (gray, PDB 5DFX) and that of miRFP670nano (red, PDB 6MGH). The truncated heptad in GAF3 is shown in blue. C) UV-visible absorption spectra of GAF3_W496Y_-PEB (blue) and trGAF3_W496Y_-PEB (red), each expressed from a single plasmid system. The region depicting bound PEB/PUB is enlarged in the inset. D) Fluorescence spectra of GAF3_W496Y_-PEB (blue) and trGAF3_W496Y_-PEB (red). Fluorescence spectra were normalized to maximum emission of each protein. λ_ex_ = 490 nm. Protein concentration = 3 μM. Buffer = 50 mM HEPES, 100 mM NaCl, pH 7.1.

Given that mutation at W496 led to significant changes in the spectroscopic properties of full-length GAF3-PEB and improved chromophorylation, we exploited the single plasmid expression system, pET28a-PebS-trGAF3_W496Y_-HO1, to carry out site saturation mutagenesis of Y496 in truncated GAF3. The brightest mutants were screened by measuring fluorescence intensity of each cell culture after induction and overnight growth. The top five brightest variants isolated and sequenced were trGAF3_W496Y_ (the template), trGAF3 (truncated version of the wildtype sequence), trGAF3_W496H_, trGAF3_W496V_, and trGAF3_W496G_. Among these, two new residues in the W496 position were found that were not investigated in full-length GAF3, valine and glycine. PyMOL models of these mutants suggest that these side chains could allow significantly more space in the bilin-binding pocket compared to the two earlier mutants, W496Y and W496H, albeit without any aromatic character and therefore no possibility of π-π stacking interactions (Figure 4A-C). Like trGAF3_W496Y_-PEB, trGAF3-PEB and trGAF3_W496H_-PEB show similar spectra to the corresponding full-length variants, although there are some minor differences in quantum yield and extinction coefficient values between the W496H mutants (Figure 1E-F, 4D-E, S5, S11A-F, and Table 1). The PEB/PUB chromophorylation of trGAF3_W496Y_, which is the only mutant that can be directly compared since both full-length and truncated proteins were expressed from the single plasmid system, is similar to GAF3_W496Y_. trGAF3 and trGAF3_W496H_ both have the highest chromophorylation measured, at ∼19% for PEB with an additional ∼4% of PUB. The two new mutants, trGAF3_W496V_-PEB and trGAF3_W496G_-PEB, have absorption and fluorescence properties and spectra that generally fall between trGAF3-PEB, trGAF3_W496Y_-PEB, and trGAF3_W496H_-PEB (Figure 4, S11). The W496V mutant had ∼12% chromophorylation with PEB and ∼3% PUB. trGAF3_W496G_ yielded similar chromophorylation to trGAF3_W496Y_ at ∼8% PEB and 1.5% PUB, which is the lowest of all of the truncated variants.

**Figure 4.**
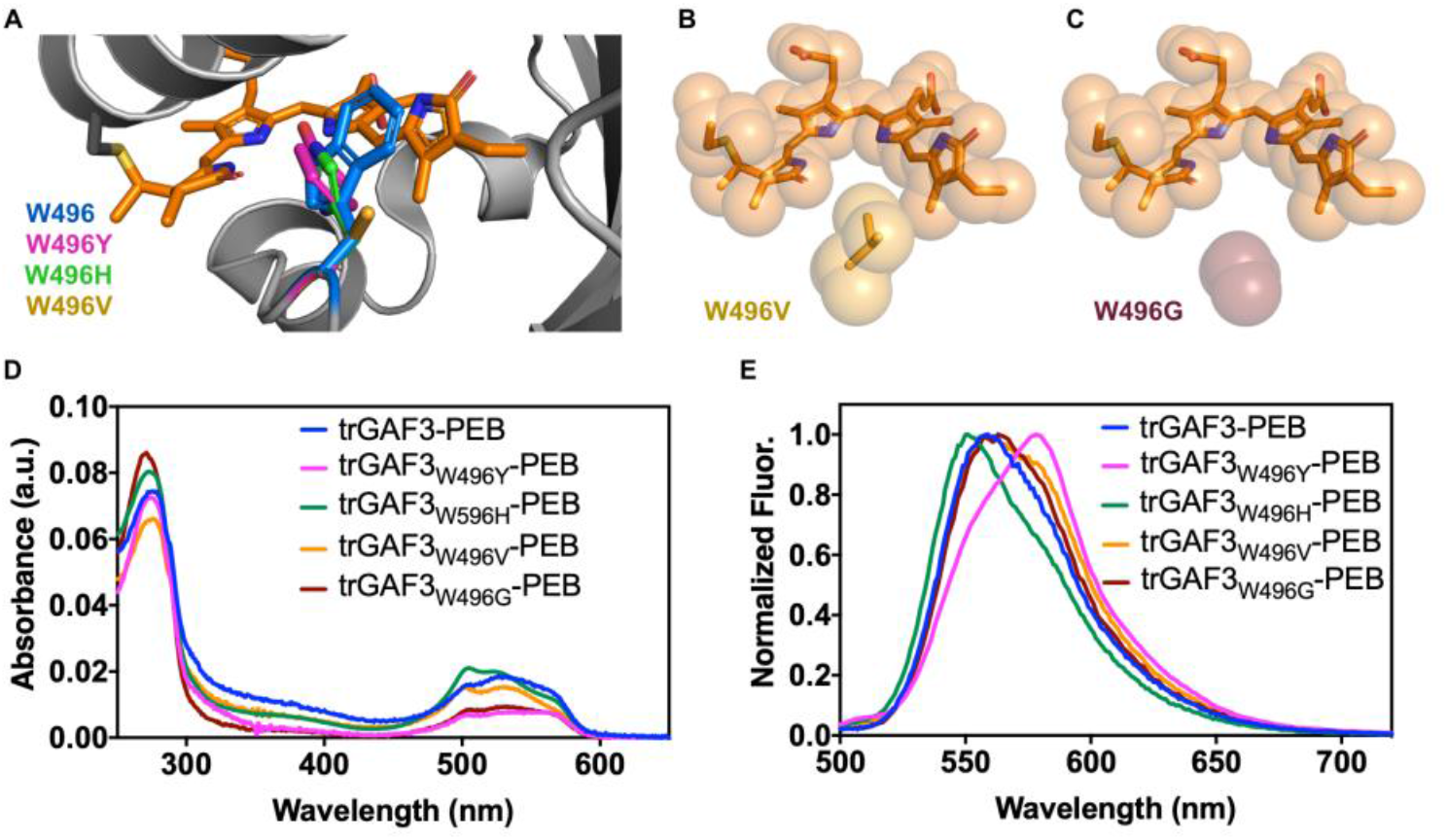
Structures and UV-visible and fluorescence spectroscopy of site saturation mutants of trGAF3_W496Y_. A) PyMOL model of the crystal structure of GAF3-PCB in the red state (PDB 5DFX). The protein structure is gray, the PCB cofactor is orange, and the Trp496 side chain is blue. Side chains for each mutant are overlaid: Trp496 in magenta, His496 in green, and Val496 in gold. Mutations were modeled using PyMOL mutagenesis wizard. B) Space-filling model of the chromophore and W496V mutant from part A. The Val side chain is gold. C) Space-filling model of the chromophore and W496G mutant from part A. Gly is colored maroon. UV-visible absorption (D) and fluorescence (E) spectra of trGAF3-PEB (blue), trGAF3_W496Y_-PEB (magenta), trGAF3_W496H_-PEB (green), trGAF3_W496V_-PEB (orange), and trGAF3_W496G_-PEB (maroon). Fluorescence spectra were normalized to maximum emission for each protein. Protein concentrations = ∼3-4 μM. λ_ex_ = 490 nm. Buffer = 50 mM HEPES, 100 mM NaCl, pH 7.1

To determine whether we could find a pattern in predicted protein-chromophore interactions among GAF domains known to bind or not bind to PEB, we aligned the GAF3 sequence with those of several GAF domains previously investigated.^15^ In prior work, several GAF domains including Slr1393 GAF3 and All2699 GAF1 and various DXCF motif-containing blue/green photoswitching GAF domains were systematically tested for PEB binding.^15^ Although binding efficiencies were not quantified, the authors detailed which domains bound PEB and estimated the extent of isomerization to PUB.^15^ Alignment of these domains did not yield a clear trend related to the W496 position in GAF3 (Figure S12). Twelve sequences were aligned, eight of which bound PEB, and most of which partially autoisomerized to PUB (with the exception of All2699g1).^15^ Of those that bound PEB, four had aromatic residues that aligned with GAF3-W496 and the other four had Val, Ile, or Gly. Among the four that did not bind PEB, the aligned residues included Lys, Ile, and Val. From this analysis, an aromatic residue may be important but is not required in this position for binding PEB. Our site saturation mutagenesis results yielding Val and Gly mutants are consistent with the presence of these residues in some of the other PEB-binding GAF domains. None of the PEB-binding GAF domains previously investigated had a Tyr natively in this position, although Phe was present in some. Although this mutant did not arise from the site-directed mutagenesis screening, the role of the hydroxyl group on Tyr in GAF3 and other PEB-binding GAF domains is an interesting topic for future studies.

To gain further insight to the role of position 496 on the chromophore properties, we also investigated the effect of pH on fluorescence emission spectra. All truncated GAF3 proteins showed pH-dependent shifts in emission spectra from pH 4.0 to 9.0 (Figure 5). When excited at 490 nm, trGAF3 and the two aromatic residue-based mutants trGAF3_W496Y_ and trGAF3_W496H_ emit with a maximum around 570 nm at low pH (4.0) that then decreases in intensity and undergoes a shift around pH 6.0. In trGAF3-PEB and trGAF3_W496H_, the emission shifts to mostly ∼560 nm with a shoulder around 580 nm, but in trGAF3-PEB_W496Y_, the shift is to a maximum around 580 nm with a shoulder of ∼560 nm that becomes more distinct as the pH increases. If the ∼560 nm emission corresponds to PUB and that at 580 nm corresponds to PEB, this result suggests that pH may also tune the ratio of PEB to PUB in GAF3 proteins. There are some differences in the pH-dependent spectra for the non-aromatic mutants compared to trGAF3 and the aromatic mutants. Most noticeably, the spectra with the highest fluorescence intensities are for pH 5.0 rather than the lowest pH 4.0 value and the fluorescence intensity mostly decreased from pH 6.0 to 9.0. Similar to trGAF3 and the aromatic mutants, trGAF3_W496V_ and trGAF3_W496G_ showed emission peaks that broadened or split to ∼560 nm and 580 nm with increasing pH, albeit with different relative intensities. Collectively, these pH-dependent data suggest that the 496 position plays a role in tuning proton interactions with the chromophore that may also affect PEB/PUB isomerization.

**Figure 5.**
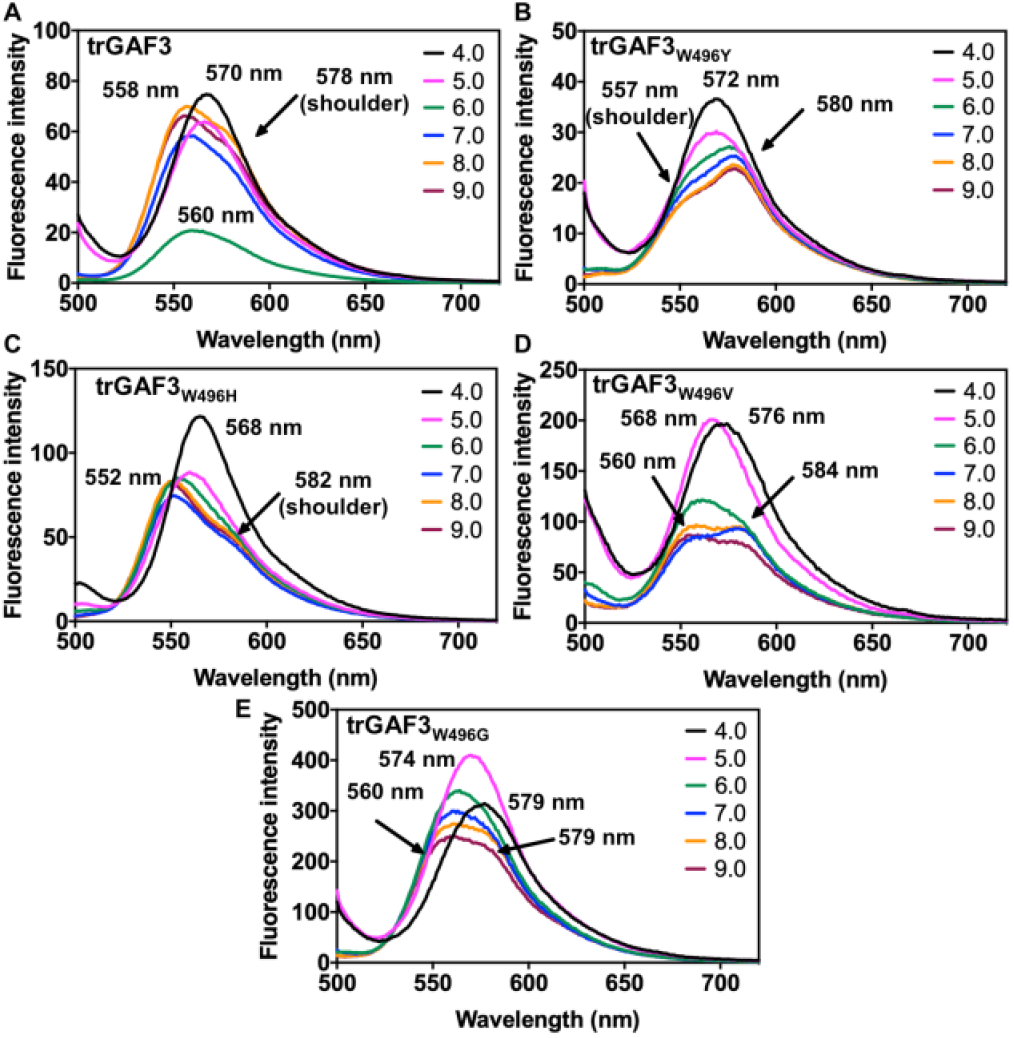
pH-dependent fluorescence emission spectra of truncated GAF3 proteins. Fluorescence emission spectra of trGAF3-PEB (A), trGAF3_W496Y_-PEB (B), trGAF3_W496H_-PEB (C), trGAF3_W496V_-PEB (D), and trGAF3_W496G_-PEB (E). Fluorescence emission spectra were collected at pH 4.0 (black), 5.0 (pink), 6.0 (green), 7.0 (blue), 8.0 (orange), and 9.0 (maroon). The most red-shifted and blue-shifted maxima and shoulders are labeled. Protein concentrations = ∼3 μM. λ_ex_ = 490 nm. Buffers: 50 mM sodium citrate, 100 mM NaCl (pH 4.0); 50 mM MES, 100 mM NaCl (pH 5.0, 6.0); 50 mM HEPES, 100 mM NaCl (pH 7.0, 8.0); 50 mM CHES, 100 mM NaCl (pH 9.0).

### Fluorescence Microscopy Imaging of GAF3 mutants

To evaluate the utility of PEB/PUB-bound GAF3 and GAF3 mutants for labeling living cells, we assessed fluorescence in *E. coli* expressing the most chromophorylated GAF3 proteins using fluorescence microscopy. Here we grew *E. coli* transformed with the single plasmid expression system and induced protein expression using the same conditions as for isolating the purified proteins with the reported chromophorylation percentages. Then, cells containing trGAF3-PEB/PUB and trGAF3_W496H_-PEB/PUB were embedded in agarose on a microscope slide and evaluated by measuring the fluorescence signal. Both truncated variants, which had up to 24% total chromophorylation when purified, showed bright fluorescence signals on the microscope (Figure 6), confirming their utility as fluorescent biomarkers.

**Figure 6.**
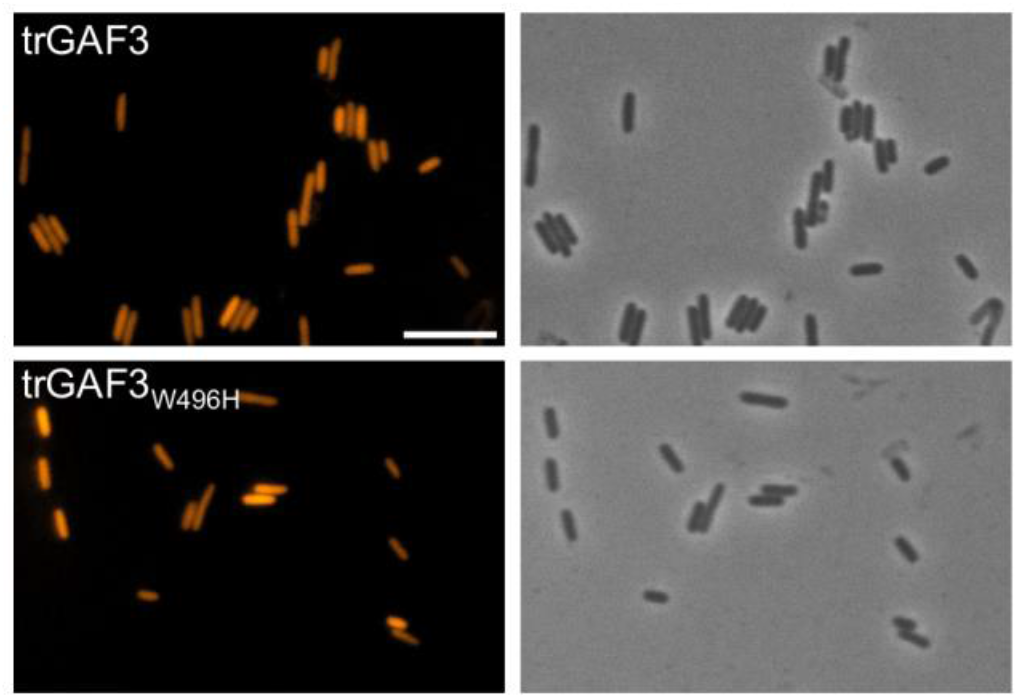
Fluorescence microscopy of *E. coli* cells expressing truncated GAF3 proteins. Top row: fluorescence signal from trGAF3 (left) and corresponding bright-field image of *E. coli* cells (right). Bottom row: fluorescence signal from trGAF3_W496H_ (left) and corresponding bright-field image of *E. coli* cells (right). Scale bar = 10 μm.

## Conclusion

In summary, we explored site-directed mutagenesis and plasmid modification routes to improve the utility of GAF3-PEB as a fluorescent biomarker in live cells. We found that mutation at a single site could more than double the percent chromophorylation of the non-native PEB cofactor, and truncation of the protein sequence further raised the chromophorylation. Plasmid modifications to adjust relative expression levels of GAF3 protein and PEB synthesis enzymes also aided in raising chromophorylation. Collectively, these efforts improved PEB loading from ∼3 to 19%. At the same time, single site mutation tuned the fluorescence emission up to ∼30 nm, likely by modifying the extent of PEB autoisomerization to the less conjugated PUB. Few studies have investigated how PEB bound to CBCRs is affected by the amino acid composition of the binding pocket. This work highlights the role of GAF3-position 496 in tuning the PEB/PUB isomerization ratio and therefore the emission wavelength over up to 30 nm. Future studies will aim to better understand protein interactions with bound PEB/PUB in CBCRs and to use these proteins as fluorescent trackers or in sensor platforms in living cells.

### Experimental Section

All reagents were purchased from commercial sources and used as received. UV-visible absorption spectra were obtained on an Agilent Cary 100 UV-visible spectrophotometer and fluorescence spectra on an Agilent Eclipse spectrofluorimeter using quartz cuvettes (Starna cells) with 1 cm path lengths.

#### Plasmid Constructs

The plasmid pSR43.6r (Addgene no. 63197) encoding heme oxygenase (HO1) and phycocyanobilin:ferredoxin oxidoreductase (PcyA) was a gift from Jeffrey Tabor.^43^ All other plasmids were constructed as described in the Supporting Information (Experimental Section, Tables S2-S6, and Figures S6-S7). All plasmids were verified by DNA sequencing. The nucleotide sequences of the promoter and coding regions of each plasmid are shown in Figures S13-S22.

#### Spectroscopic Measurements

For all spectroscopic measurements, a final concentration of 3–5 μM purified protein (expression and purification protocols in Supplemental Information) was prepared in 2.5 mL of HEPES buffer (50 mM HEPES, 100 mM NaCl, pH 7.1). Extinction coefficients for bound PEB were determined by measuring UV-visible absorbance of the folded protein over a range of concentrations and calculating the slope of the absorbance versus the concentration. The concentration of PEB-bound protein was determined by measuring the PEB (ε = 42.8 mM^-1^ cm^-1^ at 550 nm)^44^ or PUB (ε = 104 mM^-1^ cm^-1^ at 495 nm)^45^ absorbance after denaturation with acidic urea (8 M, pH 2.0). Fluorescence quantum yields (Φ) of GAF3 mutants were determined using reported Φ = 0.31 for GAF3 as a standard.^15^

#### Determination of PEB Binding Efficiency for GAF3 and Mutants

For binding efficiency measurements, absorbance spectra of 3-5 μM of protein in 2.5 mL HEPES buffer (50 mM HEPES, 100 mM NaCl, pH 7.1) were collected. The extinction coefficients of GAF3 and trGAF3 at 280 nm (εGAF3 = 34950 M^-1^ cm^-1^, εtrGAF3 = 30940 M^-1^ cm^-1^) were calculated by Expasy ProtParam.^46^ The same value was calculated for the full-length and truncated mutants. The extinction coefficient corresponding to PEB in GAF3 (εGAF3-PEB) is 43 mM^-1^ cm^-1^ (λ_max_ = 530 nm).^15^ The extinction coefficients corresponding to the absorption maxima of all mutants are reported here (Table 1). The PEB binding efficiency of GAF3-PEB was calculated by using equation 1. The PUB binding efficiency of GAF3-PUB was calculated similarly but using an absorbance maximum of 507 nm (ε = 200 mM^-1^ cm^-1^). For all other proteins, the corresponding PEB and PUB absorbance maxima along with the measure extinction coefficients were used to calculate PEB and PUB binding efficiencies (the exact values used are given in a footnote of Table S1).

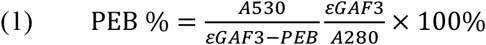

Three biological replicates (each representing a separate protein expression preparation) were used to calculate each PEB/PUB binding efficiency/chromophorylation and standard deviations (Table S1).

## Supporting Information

The Supporting Information includes an expanded Experimental Section, primers, sequence and structure alignments, additional characterizations of proteins, and DNA sequences.

## Acknowledgements

We would like to acknowledge the funding support from NIH-NIGMS grant R35 GM138223 (M.L.Z.), The Welch Foundation grant E-1972 (M.L.Z.), and the University of Houston New Faculty Startup Grant (M.L.Z.).

## Notes

The authors declare no competing financial interest.

## Notes

### Competing Interest Statement

The authors have declared no competing interest.

## References

(1) Ho, M. Y., Soulier, N. T., Canniffe, D. P., Shen, G., and Bryant, D. A. (2017) Light regulation of pigment and photosystem biosynthesis in cyanobacteria. Curr. Opin. Plant Biol. 37, 24–33.

(2) Ikeuchi, M., and Ishizuka, T. (2008) Cyanobacteriochromes: A new superfamily of tetrapyrrole-binding photoreceptors in cyanobacteria. Photochem. Photobiol. Sci. 7, 1159–1167.

(3) Rockwell, N. C., and Lagarias, J. C. (2010) A brief history of phytochromes. ChemPhysChem 11, 1172–1180.

(4) Anders, K., and Essen, L. O. (2015) The family of phytochrome-like photoreceptors: Diverse, complex and multi-colored, but very useful. Curr. Opin. Struct. Biol. 35, 7–16.

(5) Rockwell, N. C., Su, Y.-S., and Lagarias, J. C. (2006) Phytochrome Structure and Signaling Mechanisms. Annu. Rev. Plant Biol. 57, 837–858.

(6) Narikawa, R., Fukushima, Y., Ishizuka, T., Itoh, S., and Ikeuchi, M. (2008) A Novel Photoactive GAF Domain of Cyanobacteriochrome AnPixJ That Shows Reversible Green/Red Photoconversion. J. Mol. Biol. 380, 844–855.

(7) Hirose, Y., Shimada, T., Narikawa, R., Katayama, M., and Ikeuchi, M. (2008) Cyanobacteriochrome CcaS is the green light receptor that induces the expression of phycobilisome linker protein. Proc. Natl. Acad. Sci. U.S.A. 105, 9528–9533.

(8) Rockwell, N. C., Martin, S. S., and Lagarias, J. C. (2012) Red/green cyanobacteriochromes: Sensors of color and power. Biochemistry 51, 9667–9677.

(9) Ishizuka, T., Narikawa, R., Kohchi, T., Katayama, M., and Ikeuchi, M. (2007) Cyanobacteriochrome TePixJ of Thermosynechococcus elongatus harbors phycoviolobilin as a chromophore. Plant Cell Physiol. 48, 1385–1390.

(10) Narikawa, R., Nakajima, T., Aono, Y., Fushimi, K., Enomoto, G., Ni-Ni-Win Itoh, S., Sato, M., and Ikeuchi, M. (2015) A biliverdin-binding cyanobacteriochrome from the chlorophyll d–bearing cyanobacterium Acaryochloris marina. Sci. Rep. 5, 7950.

(11) Chen, Y., Zhang, J., Luo, J., Tu, J. M., Zeng, X. L., Xie, J., Zhou, M., Zhao, J. Q., Scheer, H., and Zhao, K. H. (2012) Photophysical diversity of two novel cyanobacteriochromes with phycocyanobilin chromophores: Photochemistry and dark reversion kinetics. FEBS J. 279, 40–54.

(12) Xu, X. L., Gutt, A., Mechelke, J., Raffelberg, S., Tang, K., Miao, D., Valle, L., Borsarelli, C. D., Zhao, K. H., and Gärtner, W. (2014) Combined mutagenesis and kinetics characterization of the bilin-binding GAF domain of the protein Slr1393 from the cyanobacterium Synechocystis PCC6803. ChemBioChem 15, 1190–1199.

(13) Zhang, J., Wu, X. J., Wang, Z. B., Chen, Y., Wang, X., Zhou, M., Scheer, H., and Zhao, K. H. (2010) Fused-gene approach to photoswitchable and fluorescent biliproteins. Angew. Chem. Int. Ed. 49, 5456–5458.

(14) Murphy, J. T., and Lagarias, J. C. (1997) The phytofluors: A new class of fluorescent protein probes. Curr. Biol. 7, 870–876.

(15) Sun, Y. F., Xu, J. G., Tang, K., Miao, D., Gärtner, W., Scheer, H., Zhao, K. H., and Zhou, M. (2014) Orange fluorescent proteins constructed from cyanobacteriochromes chromophorylated with phycoerythrobilin. Photochem. Photobiol. Sci. 13, 757–763.

(16) Frankenberg, N., and Lagarias, J. C. (2012) Biosynthesis and Biological Functions of Bilins. The Porphyrin Handbook: Chlorophylls and Bilins: Biosynthesis, Synthesis and Degradation. Elsevier Inc.

(17) Fairchild, C. D., Zhao, J., Zhou, J., Colson, S. E., Bryant, D. A., and Glazer, A. N. (1992) Phycocyanin alpha-subunit phycocyanobilin lyase. Proc. Natl. Acad. Sci. U.S.A. 89, 7017–7021.

(18) Zhao, K. H., Zhang, J., Tu, J. M., Böhm, S., Plöscher, M., Eichacker, L., Bubenzer, C., Scheer, H., Wang, X., and Zhou, M. (2007) Lyase activities of CpcS- and CpcT-like proteins from Nostoc PCC7120 and sequential reconstitution of binding sites of phycoerythrocyanin and phycocyanin β-subunits. J. Biol. Chem. 282, 34093–34103.

(19) Rockwell, N. C., Martin, S. S., Feoktistova, K., and Lagarias, J. C. (2011) Diverse two-cysteine photocycles in phytochromes and cyanobacteriochromes. Proc. Natl. Acad. Sci. U.S.A. 108, 11854–11859.

(20) Rockwell, N. C., Martin, S. S., and Lagarias, J. C. (2016) Identification of Cyanobacteriochromes Detecting Far-Red Light. Biochemistry 55, 3907–3919.

(21) Rockwell, N. C., Martin, S. S., and Lagarias, J. C. (2012) Mechanistic insight into the photosensory versatility of DXCF cyanobacteriochromes. Biochemistry 51, 3576–3585.

(22) Rockwell, N. C., Martin, S. S., Gulevich, A. G., and Lagarias, J. C. (2012) Phycoviolobilin formation and spectral tuning in the DXCF cyanobacteriochrome subfamily. Biochemistry 51, 1449–1463.

(23) Ishizuka, T., Kamiya, A., Suzuki, H., Narikawa, R., Noguchi, T., Kohchi, T., Inomata, K., and Ikeuchi, M. (2011) The cyanobacteriochrome, TePixJ, isomerizes its own chromophore by converting phycocyanobilin to phycoviolobilin. Biochemistry 50, 953–961.

(24) Rockwell, N. C., Moreno, M. V., Martin, S. S., and Lagarias, J. C. (2022) Protein–chromophore interactions controlling photoisomerization in red/green cyanobacteriochromes. Photochem Photobiol Sci 21, 471–491.

(25) Zhao, K.-H., Su, P., Tu, J.-M., Wang, X., Liu, H., Ploscher, M., Eichacker, L., Yang, B., Zhou, M., and Scheer, H. (2007) Phycobilin:cystein-84 biliprotein lyase, a near-universal lyase for cysteine-84-binding sites in cyanobacterial phycobiliproteins. Proc. Natl. Acad. Sci. U.S.A. 104, 14300–14305.

(26) Tang, K., Zeng, X. L., Yang, Y., Wang, Z. B., Wu, X. J., Zhou, M., Noy, D., Scheer, H., and Zhao, K. H. (2012) A minimal phycobilisome: Fusion and chromophorylation of the truncated core-membrane linker and phycocyanin. Biochim. Biophys. Acta Bioenerg. 1817, 1030–1036.

(27) Fushimi, K., Miyazaki, T., Kuwasaki, Y., Nakajima, T., Yamamoto, T., Suzuki, K., Ueda, Y., Miyake, K., Takeda, Y., Choi, J. H., Kawagishi, H., Park, E. Y., Ikeuchi, M., Sato, M., and Narikawa, R. (2019) Rational conversion of chromophore selectivity of cyanobacteriochromes to accept mammalian intrinsic biliverdin. Proc. Natl. Acad. Sci. U.S.A. 116, 8301–8309.

(28) Narikawa, R., Ishizuka, T., Muraki, N., Shiba, T., Kurisu, G., and Ikeuchi, M. (2013) Structures of cyanobacteriochromes from phototaxis regulators AnPixJ and TePixJ reveal general and specific photoconversion mechanism. Proceedings of the National Academy of Sciences 110, 918–923.

(29) Rockwell, N. C., Martin, S. S., Gulevich, A. G., and Lagarias, J. C. (2014) Conserved phenylalanine residues are required for blue-shifting of cyanobacteriochrome photoproducts. Biochemistry 53, 3118–3130.

(30) Xu, X., Höppner, A., Wiebeler, C., Zhao, K., Schapiro, I., and Gärtner, W. (2020) Structural elements regulating the photochromicity in a cyanobacteriochrome. Proc. Natl. Acad. Sci. U.S.A. 117, 2432–2440.

(31) Li, L., Murphy, J. T., and Lagarias, J. C. (1995) Continuous Fluorescence Assay of Phytochrome Assembly in Vitro. Biochemistry 34, 7923–7930.

(32) Slavov, C., Fischer, T., Barnoy, A., Shin, H., Rao, A. G., Wiebeler, C., Zeng, X., Sun, Y., Xu, Q., Gutt, A., Zhao, K.-H., Gärtner, W., Yang, X., Schapiro, I., and Wachtveitl, J. (2020) The interplay between chromophore and protein determines the extended excited state dynamics in a single-domain phytochrome. Proc. Natl. Acad. Sci. U.S.A. 117, 16356–16362.

(33) Camara-Artigas, A., Bacarizo, J., Andujar-Sanchez, M., Ortiz-Salmeron, E., Mesa-Valle, C., Cuadri, C., Martin-Garcia, J. M., Martinez-Rodriguez, S., Mazzuca-Sobczuk, T., Ibañez, M. J., and Allen, J. P. (2012) pH-dependent structural conformations of B-phycoerythrin from Porphyridium cruentum. FEBS J. 279, 3680–3691.

(34) Oliinyk, O. S., Shemetov, A. A., Pletnev, S., Shcherbakova, D. M., and Verkhusha, V. V. (2019) Smallest near-infrared fluorescent protein evolved from cyanobacteriochrome as versatile tag for spectral multiplexing. Nat. Commun. 10, 1–13.

(35) Rockwell, N. C., Martin, S. S., Gan, F., Bryant, D. A., and Lagarias, J. C. (2015) NpR3784 is the prototype for a distinctive group of red/green cyanobacteriochromes using alternative Phe residues for photoproduct tuning. Photochem. Photobiol. Sci. 14, 258–269.

(36) Jones, K. L., Kim, S.-W., and Keasling, J. D. (2000) Low-Copy Plasmids can Perform as Well as or Better Than High-Copy Plasmids for Metabolic Engineering of Bacteria. Metab. Eng. 2, 328–338.

(37) Salis, H. M., Mirsky, E. A., and Voigt, C. A. (2009) Automated design of synthetic ribosome binding sites to control protein expression. Nat. Biotechnol. 27, 946–950.

(38) Jiang, W., and Fang, B. (2016) Construction of a tunable multi-enzyme-coordinate expression system for biosynthesis of chiral drug intermediates. Sci. Rep. 6.

(39) Zhong, C., Wei, P., and Zhang, Y. P. (2017) Enhancing functional expression of codon-optimized heterologous enzymes in Escherichia coli BL21(DE3) by selective introduction of synonymous rare codons. Biotechnol. Bioeng. 114, 1054–1064.

(40) Chen, H., Huang, R., and Zhang, Y. H. P. (2017) Systematic comparison of co-expression of multiple recombinant thermophilic enzymes in Escherichia coli BL21(DE3). Appl. Microbiol. Biotechnol. 101, 4481–4493.

(41) Hörner, M., Gerhardt, K., Salavei, P., Hoess, P., Härrer, D., Kaiser, J., Tabor, J. J., and Weber, W. (2019) Production of Phytochromes by High-Cell-Density E. coli Fermentation. ACS Synth. Biol. 8, 2442–2450.

(42) Chen, H., and Jiang, P. (2019) Metabolic engineering of Escherichia coli for efficient biosynthesis of fluorescent phycobiliprotein. Microb. Cell Fact. 18, 1–13.

(43) Schmidl, S. R., Sheth, R. U., Wu, A., and Tabor, J. J. (2014) Refactoring and optimization of light-switchable Escherichia coli two-component systems. ACS Synth. Biol. 3, 820–831.

(44) Glazer, A. N., and Hixson, C. S. (1975) Characterization of R phycocyanin. Chromophore content of R phycocyanin and C phycoerythrin. J. Biol. Chem. 250, 5387–5495.

(45) Glazer, A. N., and Hixson, C. S. (1977) Subunit structure and chromophore composition of rhodophytan phycoerythrins. Porphyridium cruentum B phycoerythrin and b phycoerythrin. J. Biol. Chem. 252, 32–42.

(46) Gasteiger, E., Hoogland, C., Gattiker, A., Duvaud, S., Wilkins, M. R., Appel, R. D., and Bairoch, A. (2005) Protein Identification and Analysis Tools on the ExPASy Server, in The Proteomics Protocols Handbook, pp 571–607. Humana Press, Totowa, NJ.

