## Supporting Information for "A Single Site Mutation Tunes Fluorescence and Chromophorylation of an Orange Fluorescent Cyanobacteriochrome"

### Table of Contents


Experimental Section ........ S2

Table S1. Summary of chromophorylation ratios ........ S6

Table S2. Primer sequences for GAF3 Trp496 site mutations and varied RBS sequences ........ S6

Table S3. Primer sequences for pACYCDuet-HO1-PcyA and pACYCDuet-PebS-HO1 ........ S7

Table S4. Primer sequences for pET28a-PebS-HO1 and pACYCDuet-GAF3$_{W496Y}$ ........ S7

Table S5. Primer sequences for pACYCDuet-PebS-T7-ter-HO1 and pET28a-PebS-GAF3$_{W496Y}$-HO1 ........ S8

Table S6. Primer sequences for GAF3$_{W496Y}$ truncation and site saturation mutagenesis ........ S8

Scheme S1. Biosynthesis and covalent binding of phycocyanobilin (PCB) and phycoerythrobilin (PEB) to GAF3 ........ S9

Figure S1. SDS-PAGE of GAF3 proteins ........ S10

Figure S2. UV-visible absorption and fluorescence spectra of GAF3-PCB ........ S11

Figure S3. Crystal structure overlays of Slr1393 GAF3-PCB with PEB-bound proteins and Cph1-PCB ........ S12

Figure S4. Multiple sequence alignment of Slr1393 GAF3 with other CBCRs and the GAF domains of phytochromes ........ S13

Figure S5. Fluorescence excitation and emission spectra of GAF3 and GAF3 mutants ........ S14

Figure S6. Schematic of PCR amplification to generate the PebS-T7 ter fragment ........ S15

Figure S7. Schematic of PCR amplification to generate the PebS-T7 ter-HO1 fragment ........ S16

Figure S8. UV-visible absorption spectra depicting chromophorylation of GAF3$_{W496Y}$ from pACYCDuet vs pET28a ........ S17

Figure S9. Plasmid designs and UV-visible absorption spectra for GAF3$_{W496Y}$ expression with varied RBS sequences ........ S18

Figure S10. UV-visible absorption spectra depicting chromophorylation of GAF3$_{W496Y}$ from single vs dual plasmid ........ S19

Figure S11. Fluorescence excitation and emission spectra of truncated GAF3 and GAF3 mutants ........ S20

Figure S12. Multiple sequence alignment of Slr1393 GAF3 with other CBCRs tested for PEB binding ........ S21

Figure S13. Nucleotide sequence of the promoter and coding regions of pET28a-GAF3 ........ S22

Figure S14. Nucleotide sequence of the promoter and coding regions of pET28a-GAF3$_{W496Y}$ ........ S23

Figure S15. Nucleotide sequence of the promoter and coding regions of pET28a-GAF3$_{W496H}$ ........ S24

Figure S16. Nucleotide sequence of the promoter and coding regions of pACYCDuet-HO1-PcyA ........ S25

Figure S17. Nucleotide sequence of the promoter and coding regions of pACYCDuet-PebS-HO1 ........ S26

Figure S18. Nucleotide sequence of the promoter and coding regions of pET28a-PebS-HO1 ........ S27

Figure S19. Nucleotide sequence of the promoter and coding regions of pACYCDuet-GAF3$_{W496Y}$ ........ S28

Figure S20. Nucleotide sequence of the promoter and coding regions of pACYCDuet-PebS-T7-ter-HO1 ........ S29

Figure S21. Nucleotide sequence of the promoter and coding regions of pET28a-PebS-GAF3$_{W496Y}$-HO1 ........ S30

Figure S22. Nucleotide sequence of the promoter and coding regions of pET28a-PebS-trGAF3$_{W496Y}$-HO1 ........ S31

### Experimental Section

#### General Information

Primers were ordered from Sigma-Aldrich. *E. coli* strains [DH5α and BL21(DE3)] for cloning and recombinant protein expression were obtained from New England Biolabs. NEBuilder HiFi DNA Assembly, restriction enzymes, Q5 DNA polymerase, T4 DNA ligase, and DNAse I were also obtained from New England Biolabs. PCR cleanup and gel extraction kits were from New England Biolabs. Plasmid DNA miniprep kits were from New England Biolabs or Zymo Research. Columns for purification were from Bio-Rad. Protein gel electrophoresis was carried out using Bio-Rad 4–20% Mini-PROTEAN TGX Stain-Free Protein Gels and a mini gel apparatus. Gels were imaged on a Bio-Rad ChemiDoc Touch Imaging System. Fast Protein Liquid Chromatography was carried out on a Bio-Rad NGC Quest 10 Chromatography system.

#### Plasmid Constructs

The plasmid pSR43.6r (Addgene no. 63197) encoding heme oxygenase (HO1) and phycocyanobilin:ferredoxin oxidoreductase (PcyA) was a gift from Jeffrey Tabor.[1] All plasmids were verified by DNA sequencing.

***pET28a-GAF3.*** The gene encoding for the GAF3 domain from Slr1393 of *Synechocystis* PCC 6803 (NCBI protein ID BAA17210, amino acids L441–Q596) with a TGA stop codon and inserted between BamHI and XhoI restriction sites in pET28a was ordered from Genscript. The nucleotide sequence of the promoter and coding regions is shown in Figure S13.

***pET28a-GAF3$_{W496Y}$* and *pET28a-GAF3$_{W496H}$*.** The plasmids encoding for GAF3 mutants were prepared by site-directed PCR mutagenesis (Q5 site-directed mutagenesis, NEB) of pET28a-GAF3 using the primers listed in Table S2. The nucleotide sequences of the promoter and coding regions is shown in Figures S14-S15.

***pMH48-GAF3$_{W496Y}$*, *pMH51-GAF3$_{W496Y}$*, and *pMH52-GAF3$_{W496Y}$*.** The plasmids encoding for varied ribosome binding sites were prepared by site-directed PCR mutagenesis (Q5 site-directed mutagenesis, NEB) of pET28a-GAF3$_{W496Y}$ with the primers listed in Table S2.

***pACYCDuet-HO1-PcyA.*** The plasmid for the co-expression of heme oxygenase (HO1) and phycocyanobilin:ferredoxin oxidoreductase (PcyA) was cloned using pSR43.6r (Addgene no. 63197) and the plasmid pACYCDuet-1 (Novagen). The *ho1-pcya* gene fragment was PCR amplified from pSR43.6r using the primers listed in Table S3 and inserted between EcoRI and SacI restriction sites in pACYCDuet-1. The nucleotide sequence of the promoter and coding regions is shown in Figure S16.

***pACYCDuet-PebS-HO1.*** The initial plasmid for the co-expression of HO1 and phycoerythrobilin synthase (PebS) was cloned using the gene for PebS from *Prochlorococcus phage P-SSM2* ordered from Genscript in the pUC18 plasmid (codon optimized for *E. coli*), the plasmid pSR43.6r (Addgene no. 63197), and the plasmid pACYCDuet-1 (Novagen). The *ho1* gene fragment was PCR amplified from pSR43.6r using the primers listed below and inserted between NdeI and XhoI restriction sites in pACYCDuet-1. The *pebs* gene fragment from pUC18-PebS (Genscript) was PCR-amplified and assembled with PCR-amplified pACYCDuet-ho1 by NEBuilder HiFi DNA Assembly using the primers listed in Table S3. The nucleotide sequence of the promoter and coding regions is shown in Figure S17.

***pET28a-PebS-HO1.*** The co-expression plasmid for HO1 and PebS enzymes with an additional T7 promoter and T7 terminator between the PebS and HO1 in pET28a vector was prepared using pACYCDuet-PebS-HO1 and pET28a-GAF3. First, the *pebs* gene fragment was PCR-amplified from pACYCDuet-PebS-HO1, and the T7 terminator (*T7ter*) gene was PCR-amplified from pET28a-GAF3. Then, the *PebS-T7ter* fragment was ligated by overlap extension PCR (Figure S6). The *ho1* gene fragment including T7 promoter and T7 terminator was PCR-amplified from pACYCDuet-PebS-HO1. Then, the PebS-T7ter-HO1 fragment was ligated by overlap extension PCR (Figure S7) and assembled with PCR-amplified pET28a vector by NEBuilder HiFi DNA Assembly. All primers are listed in Table S4. The nucleotide sequence of the promoter and coding regions is shown in Figure S18.

***pACYCDuet-GAF3$_{W496Y}$*.** The GAF3$_{W496Y}$ fragment including the N-terminal $His_6$ tag, thrombin site, and T7 tag, and the C-terminal T7 terminator was PCR-amplified from pET28a-GAF3$_{W496Y}$ using the primers listed in Table S4. The PCR product was inserted between NcoI and SacI restriction sites in pACYCDuet-1 (Novagen). The nucleotide sequence of the promoter and coding regions is shown in Figure S19.

***pACYCDuet-PebS-T7-ter-HO1.*** The PebS-T7 ter-HO1 fragment (from cloning of pET28a-PebS-HO1 described above) was PCR-amplified and assembled with PCR-amplified pACYCDuet-1 vector by NEBuilder HiFi DNA Assembly using the primers listed in Table S5. The nucleotide sequence of the promoter and coding regions is shown in Figure S20.

***pET28a-PebS-GAF3$_{W496Y}$-HO1.*** The GAF3$_{W496Y}$ fragment was PCR-amplified from pET28a-GAF3$_{W496Y}$ and inserted between PebS and HO1 in the PCR-amplified pET28a-PebS-HO1 plasmid with NEBuilder HiFi DNA Assembly using the primers listed in Table S5. The nucleotide sequence of the promoter and coding regions is shown in Figure S21.

***pET28a-PebS-trGAF3$_{W496Y}$-HO1.*** Truncation of the protein was achieved by performing PCR on pET28a-PebS-GAF3$_{W496Y}$-HO1 using Q5 Hot Start High-Fidelity 2X Master Mix (NEB) with the primers listed in Table S6. The PCR product was treated with KLD enzyme mix (NEB) prior to transformation in DH5α *E. coli* cells (NEB). The nucleotide sequence of the promoter and coding regions is shown in Figure S22.

**Protein Expression and Purification**

All plasmids were (co)transformed in *E. coli* BL21(DE3) cells and selected by dual antibiotics kanamycin (50 µg/mL) and chloramphenicol (25 µg/mL) on an LB/agar plate (kanamycin only for the single plasmid expression system).

For GAF3-PCB co-expression, LB medium (100 mL) containing antibiotics was inoculated with bacterial colonies and grown at 37 °C, 250 rpm overnight. An aliquot of this starter culture (10 mL) was used to inoculate kanamycin- and chloramphenicol-containing LB broth (1 L in a 2-liter baffled flask) to an $OD_{600}$ of 0.04. After growth at 37 °C and 250 rpm for 2 hours to an $OD_{600}$ of 0.4−0.6, co-expression was induced by 0.5 mM IPTG. The induced culture was then incubated with shaking (200 rpm) for 16 h at 18 °C in the dark, and then the cells were harvested for protein purification.[2]

For full-length and truncated GAF3-PEB and GAF3 mutants-PEB (co-)expression, LB medium (100 mL) containing 0.5% w/v glucose and antibiotics was inoculated with bacterial colonies and grown at 37 °C, 250 rpm overnight. An aliquot of this starter culture (20 mL) was used to inoculate 400 mL kanamycin- and chloramphenicol-containing terrific broth (TB) in a 2-liter unbaffled flask to an $OD_{600}$ of 0.1. After growing the culture at 37 °C and 250 rpm for 90 minutes to an $OD_{600}$ of 0.6−0.8, 0.5% w/v glucose was added and (co-)expression was induced by 0.5 mM IPTG. The induced culture was further incubated for 24 h at 18 °C with shaking at 120 rpm in the dark, and then the cells were harvested for protein purification.[3]

GAF3-PCB and full length or truncated GAF3 (or GAF3$_{W496Y}$, GAF3$_{W496H}$)-PEB were FPLC-purified via immobilized metal affinity chromatography (IMAC). For His tag removal, the eluted protein was exchanged with thrombin cleavage buffer (50 mM Tris, 100 mM NaCl, 2.5 mM $CaCl_2$, 0.1% β-mercaptoethanol, pH 8.0) and concentrated to 2 mL using an Amicon Ultra 10-kDa MWCO centrifugal filter. The N-terminal $His_6$-tag from the protein was cleaved by treatment with thrombin protease overnight at 4 °C (thrombin:protein fusion ratio of 1:500 (w/w), protein concentration was determined using the Bradford Assay). Then the sample was repurified by IMAC and the purified protein was concentrated, then 10% (v/v) glycerol was added before flash freezing with liquid nitrogen and storing at –80°C. Storage was done in small aliquots (~100 µL) to prevent protein aggregation as a result of repetitive freeze-thaw cycles. Protein fractions were assayed for homogeneity by denaturing polyacrylamide gel electrophoresis (SDS-PAGE, Figure S1) and concentrations determined using the Bradford Assay or absorbance at 280 nm.

### Site Saturation Mutagenesis

Site saturation mutagenesis was accomplished by PCR on pET-ho1-pebs-trGAF3$_{W496Y}$ using the Q5 Site-Directed Mutagenesis Kit (NEB) and a degenerate nucleotide, NNK. The targeted codon for position 496 was replaced with NNK where N is any nucleotide and K is G or T, thereby coding for all 20 amino acids. The primers used are listed in Table S6. The PCR products contain the degenerate NNK substitution at the codon for position 496. After PCR the mixture was treated with KLD (kinase ligase DpnI) enzyme mix (NEB) and then transformed into BL21(DE3) *E. coli* cells (NEB) by heat shock. Transformants were selected by kanamycin resistance.

### Screening of Truncated Mutants

Single colony transformants from the site saturation mutagenesis library were inoculated into individual wells of a 96-well plate containing 200 µL of RM + glucose medium (1X M9 salts, 2% casamino acids, 0.2% glucose, 1 mM $MgCl_2$, kanamycin 30 µg/mL). The cultures were grown at 37 °C with shaking at 150 rpm for 3-4 hours in a humidity chamber. The cultures were then induced with IPTG (1 mM) and further grown overnight at 20 °C. The fluorescence properties of the mutants were screened using a Tecan plate reader with $\lambda_{ex}$ = 520 nm and slit widths = 10 nm (ex), 10 nm (em). The mutants with the highest fluorescence intensities were selected, and the corresponding cultures from the microtiter plate were used to inoculate 5 mL of fresh LB media (50 µg/mL kanamycin). These inoculates were grown overnight at 37 °C and then the plasmids were isolated by miniprep (NEB) for DNA sequencing.

### Assessment of UV-Visible Absorption and Fluorescence Spectra with Varied pH

The fluorescence of truncated GAF3 and truncated GAF3 mutants (3 µM) were measured at various pH values ranging from 4.0-9.0. The emission spectra were recorded with excitation at 490 nm. Buffers used were as follows: 50 mM sodium acetate, 100 mM NaCl for pH 4.0; 50 mM MES, 100 mM NaCl for pH 5.0 and 6.0; 50 mM HEPES for pH 7.0 and 8.0, 100 mM NaCl; 50 mM CHES, 100 mM NaCl for pH 9.0.

### Fluorescence Microscopy of Truncated GAF3 Proteins

Similar to the protein expression protocol, single colony *E. coli* transformants were grown in LB medium with antibiotics overnight prior to dilution in TB with antibiotics and growth to $OD_{600}$ of 0.7-0.8. After IPTG induction (0.5 mM), cells were incubated overnight at 20 °C. Cells were centrifuged, washed with 50 mM HEPES, 100 mM NaCl, pH 7.1 and resuspended in buffer. The cell suspension was applied to a tenth of a millimeter thick 3% agarose pad, which was sandwiched between a glass cover slip and microscope slide. Images were collected using an ECHO Revolve microscope equipped with a 5 MP CMOS Monochrome Camera (fluorescence) and a 12 MP Color Camera. The light source was a LED light cube-FITC(M) (ex:470/40, em: 525/50). Images were obtained with an oil-immersion objective at 100× magnification.

**Table S1.** PCB or PEB Binding Efficiency of GAF3 or GAF3 Mutants.

| Plasmid | GAF3/mutants | Chromophore | Chromophorylation (%)[a] |
|---|---|---|---|
| pET28a | GAF3 | PCB | 61% |
| pET28a | GAF3 | PEB/PUB | 3.0 ± 0.7 % (PEB) / 0.6 ± 0.1 % (PUB) |
| pET28a | $GAF3_{W496Y}$ | PEB/PUB | 6 ± 2 % (PEB) / 1.3 ± 0.4 % (PUB) |
| pET28a | $GAF3_{W496H}$ | PEB/PUB | 5 ± 1 % (PEB) / 11 ± 2 % (PUB) |
| pACYCDuet | $GAF3_{W496Y}$ | PEB/PUB | 11 ± 2 % (PEB) / 2.0 ± 0.5 % (PUB) |
| pMH51 | $GAF3_{W496Y}$ | PEB/PUB | 9 ± 2 % (PEB) / 1.8 ± 0.3 % (PUB) |
| pMH52 | $GAF3_{W496Y}$ | PEB/PUB | 4 ± 2 % (PEB) / 0.9 ± 0.5 % (PUB) |
| pET28a-single[b] | $GAF3_{W496Y}$ | PEB/PUB | 7.2 ± 1.6 % (PEB) / 1.6 ± 0.4 % (PUB) |
| pET28a-single[b] | $trGAF3_{W496Y}$ | PEB/PUB | 9 ± 2 % (PEB) / 2.0 ± 0.5 % (PUB) |
| pET28a-single[b] | trGAF3 | PEB/PUB | 19 ± 4 % (PEB) / 4.1 ± 0.9 % (PUB) |
| pET28a-single[b] | $trGAF3_{W496H}$ | PEB/PUB | 19 ± 6 % (PEB) / 4 ± 1 % (PUB) |
| pET28a-single[b] | $trGAF3_{W496V}$ | PEB/PUB | 12 ± 3 % (PEB) / 3.1 ± 0.8 % (PUB) |
| pET28a-single[b] | $trGAF3_{W496G}$ | PEB/PUB | 8 ± 3 % (PEB) / 1.5 ± 0.6 % (PUB) |

[a] Chromophorylation of GAF3-PEB calculated using maximal $A_{530}$ and reported $\varepsilon_{530}$ =42.8 $mM^{-1}cm^{-1}$ from reference [4]. Chromophorylation of GAF3 with PUB was calculated using maximal $A_{507}$ and measured $\varepsilon_{507}$ = 200 $mM^{-1}cm^{-1}$ (Table 1). For all $GAF3_{W496Y}$-PEB, chromophorylation percentage for PEB was calculated using maximal $A_{560}$ and measured $\varepsilon_{560}$ = 46 $mM^{-1}cm^{-1}$ and for PUB was calculated using maximal $A_{505}$ and measured $\varepsilon_{505}$ = 200 $mM^{-1}cm^{-1}$ (Table 1); For the other proteins, the corresponding maximal absorbance and ε (Table 1) used for each chromophorylation percentage calculation are as follows: $GAF3_{W496H}$, PEB: $A_{522}$ and $\varepsilon_{522}$ of 44 $mM^{-1}cm^{-1}$, PUB: $A_{502}$ and $\varepsilon_{502}$ of 140 $mM^{-1}cm^{-1}$; $trGAF3_{W496Y}$, PEB: $A_{560}$ and $\varepsilon_{560}$ of 44 $mM^{-1}cm^{-1}$, PUB: $A_{505}$ and $\varepsilon_{505}$ of 180 $mM^{-1}cm^{-1}$; trGAF3, PEB: $A_{530}$ and $\varepsilon_{530}$ of 37 $mM^{-1}cm^{-1}$, PUB: $A_{506}$ and $\varepsilon_{506}$ of 151 $mM^{-1}cm^{-1}$; $trGAF3_{W496H}$, PEB: $A_{522}$ and $\varepsilon_{522}$ of 46 $mM^{-1}cm^{-1}$, PUB: $A_{505}$ and $\varepsilon_{505}$ of 205 $mM^{-1}cm^{-1}$; $trGAF3_{W496V}$, PEB: $A_{532}$ and $\varepsilon_{532}$ of 53 $mM^{-1}cm^{-1}$, PUB: $A_{502}$ and $\varepsilon_{502}$ of 214 $mM^{-1}cm^{-1}$; $trGAF3_{W496G}$, PEB: $A_{532}$ and $\varepsilon_{532}$ of 53 $mM^{-1}cm^{-1}$, PUB: $A_{506}$ and $\varepsilon_{506}$ = 237 $mM^{-1}cm^{-1}$; Calculation details are provided in the Methods section of the main text. All error bars represent standard deviation for ≥ 3 biological replicates. [b] This pET28a plasmid also contains the *ho1* and *pebs* genes (pET28a-PebS-$GAF3_{W496Y}$-HO1, Figure S14).

**Table S2.** Primer Sequences for GAF3 Trp496 Site Mutations and Varied RBS Sequences.

| Primers | Sequence |
|---|---|
| pET28a-$GAF3_{W496Y}$_For | 5'-TTTCCCCCTCTATCAAGACACCTATC-3' |
| pET28a-$GAF3_{W496Y}$_Rev | 5'-TTATTTTGCAAGTCTTTCAGTG-3' |
| pET28a-$GAF3_{W496H}$_For | 5'-TTTCCCCCTCCATCAAGACACCTATC-3' |
| pET28a-$GAF3_{W496H}$_Rev | 5'-TTATTTTGCAAGTCTTTCAG-3' |
| pMH48-$GAF3_{W496Y}$_For | 5'-GAATACATGGGCAGCAGCCATC-3' |
| pMH48-$GAF3_{W496Y}$_Rev | 5'-CTTCTGCTTAAAGTTAAACAAAATTATTTCTAGAGGGG-3' |
| pMH51-$GAF3_{W496Y}$_For | 5'-TTATCCATGGGCAGCAGCCATC-3' |
| pMH51-$GAF3_{W496Y}$_Rev | 5'-CTCCTTCTTAAAGTTAAACAAAATTATTTCTAGAGGGG-3' |
| pMH52-$GAF3_{W496Y}$_For | 5'-GAATACATGGGCAGCAGCCATC-3' |
| pMH52-$GAF3_{W496Y}$_Rev | 5'-CTCCCCCTTAAAGTTAAACAAAATTATTTCTAGAGGGG-3' |

**Table S3.** Primer Sequences for pACYCDuet-HO1-PcyA and pACYCDuet-PebS-HO1.

| Primers | Sequence |
|---|---|
| pACYCDuet-HO1-PcyA_For | 5'-GCGGCGGAATTCCATGAGTGTCAACTTAGC-3' |
| pACYCDuet-HO1-PcyA_Rev | 5'-GTCTAGAGAGCTCTTATTGGATAACATCAAATAAG ACTTG-3' |
| pACYCDuet-PebS-HO1-ho1frag_For | 5'-GCGGAACATATGGGCATGAGTGTCAACTTAGC-3' |
| pACYCDuet-PebS-HO1-ho1frag_Rev | 5'-GCTGCACTCGAGCTAGCCTTCGGAGGTG-3' |
| pACYCDuet-PebS-HO1-pebsfrag_For | 5'-CCATGGCAGGATCCATGACAAAGAACCC-3' |
| pACYCDuet-PebS-HO1-pebsfrag_Rev | 5'-GCCGAGCTCTTACTTGTAAGAGAACAG-3' |
| pACYCDuet-PebS-HO1-vector_For | 5'-TACAAGTAAGAGCTCGGCGCG-3' |
| pACYCDuet-PebS-HO1-vector_For | 5'-ATGGATCCTGCCATGGTATATCTCCTTATTAAAGTTAAACAAAATTATTT CTAC-3' |

**Table S4. Primer Sequences for pET28a-PebS-HO1 and pACYCDuet-GAF3$_{W496Y}$.**

| Primers | Sequence |
|---|---|
| pET28a-PebS-HO1-pebsfrag_For | 5'-GGAGATATACCATGACAAAGAACCCACGC-3' |
| pET28a-PebS-HO1-pebsfrag_Rev | 5'-AGCCGGATCTTACTTGTAAGAGAACAGGAAATCATT AACAAACGC-3' |
| pET28a-PebS-HO1-T7terfrag_For | 5'-ACAAGTAAGATCCGGCTGCTAACAAAGC-3' |
| pET28a-PebS-HO1-T7terfrag_Rev | 5'-CCGAGCTCCAAAAAACCCCTCAAGACCCGT-3' |
| pET28a-PebS-HO1-ho1frag_For | 5'-GTTTTTTGGAGCTCGGCGCGC-3' |
| pET28a-PebS-HO1-ho1frag_Rev | 5'-GCCGGATCTTAGCCTTCGGAGGTGGCG-3' |
| pET28a-PebS-HO1-vector_For | 5'-AGGCTAAGATCCGGCTGCTAACAAAGC-3' |
| pET28a-PebS-HO1-vector_Rev | 5'-TTCTTTGTCATGGTATATCTCCTTCTTAAAGTTAAACAAAATTATTTCTA GAGG-3' |
| pACYCDuet-GAF3$_{W496Y}$_For | 5'-GCTAATCCATGGGCAGCAGCCATC-3' |
| pACYCDuet-GAF3$_{W496Y}$_Rev | 5'-GTTCTGAGCTCCAAAAAACCCCTCAAGACCCG-3' |

**Table S5.** Primer Sequences for pACYCDuet-PebS-T7-ter-HO1 and pET28a-PebS-GAF3$_{W496Y}$-HO1.

| Primers | Sequence |
|---|---|
| pACYCDuet-PebS-T7-ter-HO1-pebsT7terho1frag_For | 5'-GTTTAACTTTAATAAGGAGATATACCATGACAAAGAACCCAC-3' |
| pACYCDuet-PebS-T7-ter-HO1-pebsT7terho1frag_Rev | 5'-AGACTCGAGTTAGCCTTCGGAGGTGGCG-3' |
| pACYCDuet-PebS-T7-ter-HO1-vector_For | 5'-GAAGGCTAACTCGAGTCTGGTAAAGAAACCG-3' |
| pACYCDuet-PebS-T7-ter-HO1-vector_Rev | 5'-TGGTATATCTCCTTATTAAAGTTAAACAAAATTATTTCTAC AGGG-3' |
| pET28a-PebS-GAF3$_{W496Y}$-HO1-gaffrag_For | 5'-CAAGCTTATGCCGGCCACGATGC-3' |
| pET28a-PebS-GAF3$_{W496Y}$-HO1-gaffrag_Rev | 5'-TTAAGCATTATGATCCGGATATAGT TCCTCCTTTCAGC-3' |
| pET28a-PebS-GAF3$_{W496Y}$-HO1-vectorpebsho1frag_For | 5'-TATATCCGGATCATAATGCTTAAGTCGAACAGAAAGTAA TCGTATTG-3' |
| pET28a-PebS-GAF3$_{W496Y}$-HO1-vectorpebsho1frag_Rev | 5'-GCCGGCATAAGCTTGTCGACCTG CAGG-3' |

**Table S6.** Primer Sequences for Truncated GAF3$_{W496Y}$ and Site Saturation Mutagenesis

| Primers | Sequence |
|---|---|
| pET28a-PebS-T7-ter-HO1-trGAF3$_{W496Y}$_For | 5'- CCTGCAGGTCGACAAGCTT -3' |
| pET28a-PebS-T7-ter-HO1-trGAF3$_{W496Y}$_Rev | 5'- GATTATGCGGCCGTGTACAATACG -3' |
| pET28a-PebS-T7-ter-HO1-trGAF3$_{W496X}$_For | 5'- TTTCCCCCTCNNKCAAGACACCTATC-3' |
| pET28a-PebS-T7-ter-HO1-trGAF3$_{W496X}$_Rev | 5'- GTGGTGCTCGAGTCAGCTTTGGTACACTG -3' |

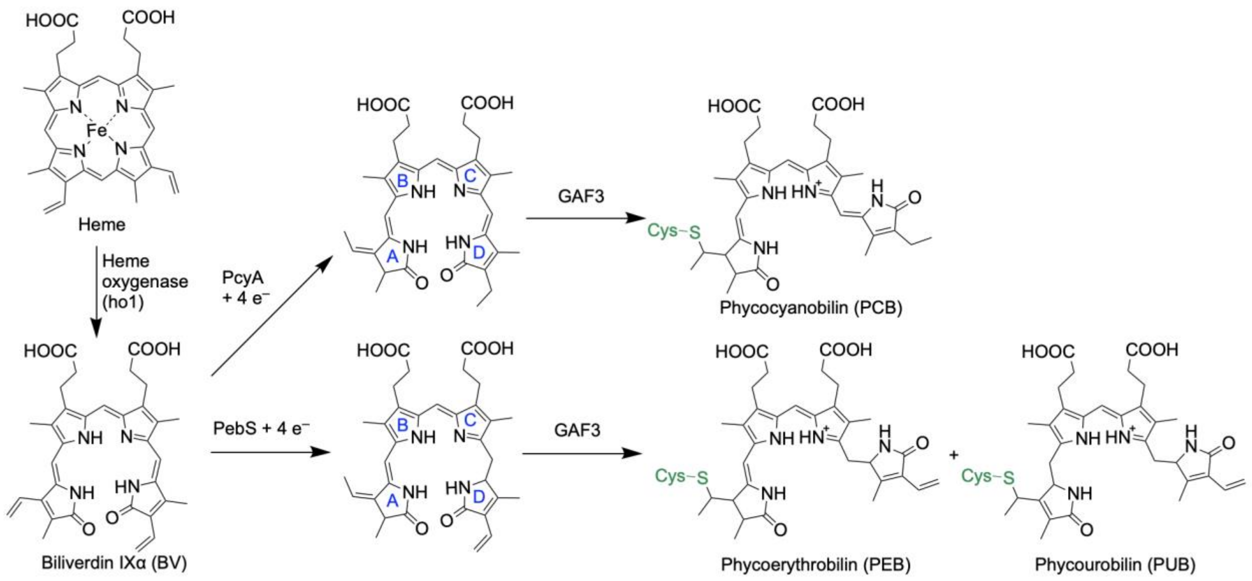


**Scheme S1.** Biosynthesis and covalent binding of phycocyanobilin (PCB) and phycoerythrobilin (PEB) to GAF3. Some PEB autoisomerizes to phycourobilin (PUB) upon binding to GAF3. Pyrrole rings A-D are as labeled on PCB and PEB.

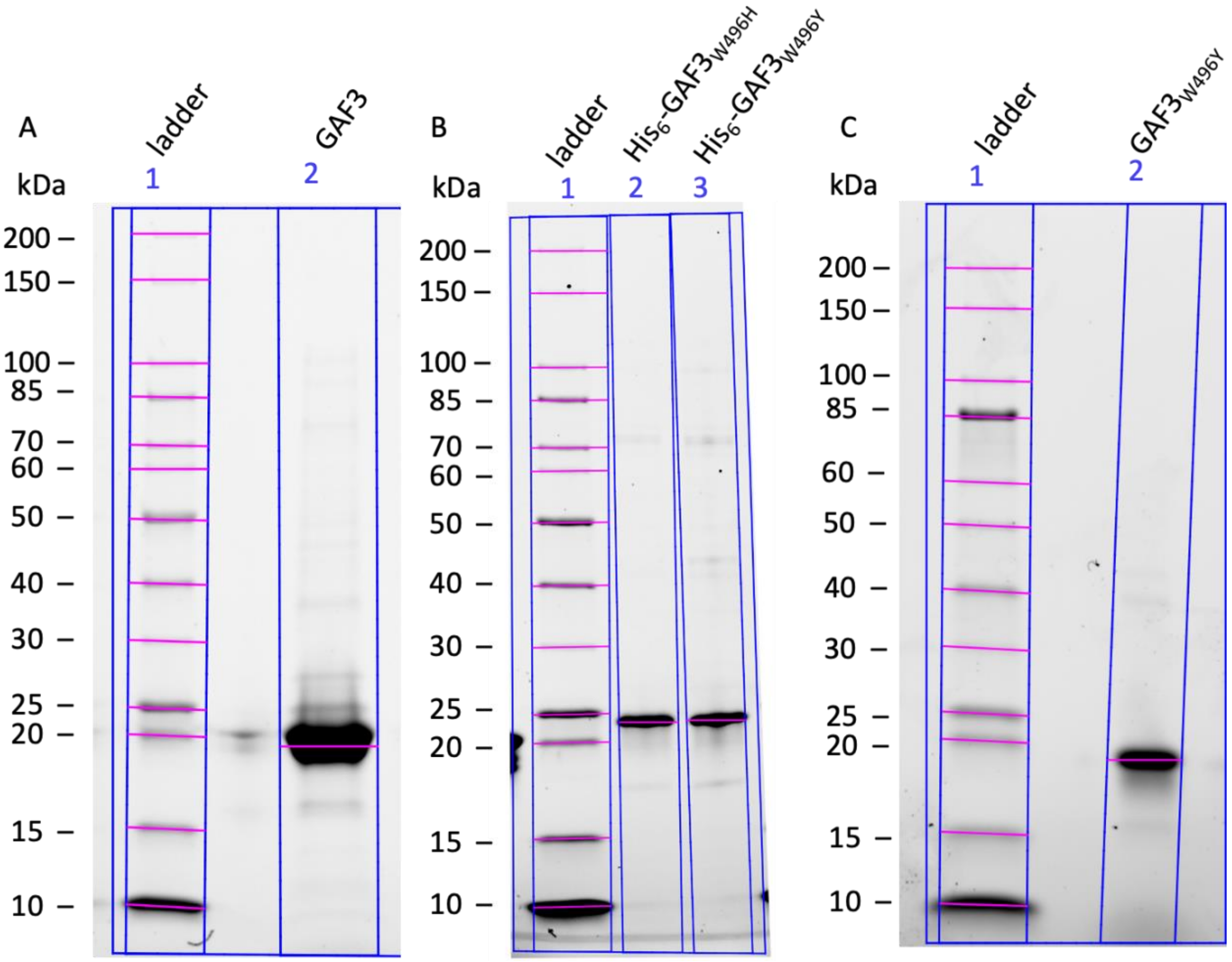


**Figure S1.** SDS-PAGE of GAF3 proteins. Denaturing polyacrylamide gel electrophoresis of purified A) GAF3, B) His$_6$-GAF3$_{W496H}$ and His$_6$-GAF3$_{W496Y}$, and C) GAF3$_{W496Y}$. Lane 1: Protein markers (molecular weights indicated in kDa), Lane 2: GAF3 (A), His$_6$-GAF3$_{W496H}$ (B), or GAF3$_{W496Y}$ (C). Lane 3: His$_6$-GAF3$_{W496Y}$ (B). Expected molecular weights are: 19.9 kDa for GAF3 and GAF3$_{W496Y}$ and 21.7 kDa for His$_6$-GAF3$_{W496H}$ and His$_6$-GAF3$_{W496Y}$. Ladder: New England Biolabs Unstained Protein Standard, Broad Range, 10–200 kDa. Gel: BioRad 4–20% Mini-PROTEAN TGX Stain-Free Protein Gels.

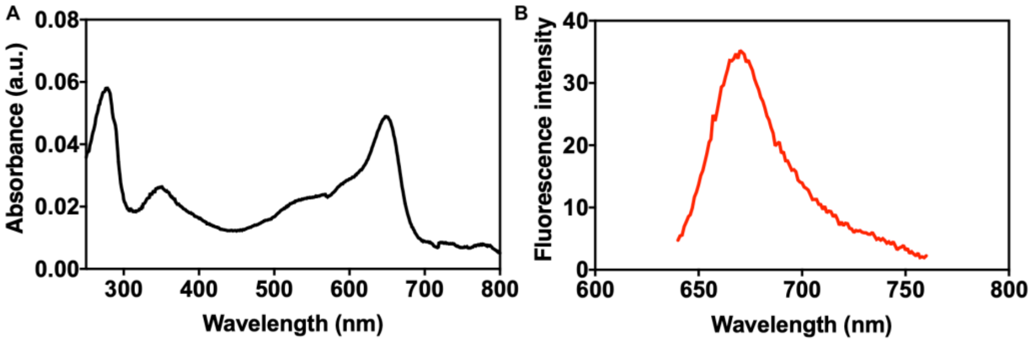


**Figure S2.** UV-visible absorption and fluorescence spectra of GAF3-PCB as isolated (mixture of red and green states). A) UV-visible absorption spectrum of GAF3-PCB (1.7 µM) and B) fluorescence spectrum of GAF3-PCB obtained upon excitation at 620 nm (red, or Z, state) with emission collected from 640-760 nm. The PCB binding efficiency was calculated by accounting for both red (A650 = 0.049, ε = 93 $mM^{-1}cm^{-1}$) and green (A539 = 0.022, ε = 50 $mM^{-1}cm^{-1}$) states in the absorption spectrum and comparing to the total protein absorbance at 280 nm (A280 = 0.057, ε = 34950).[2] The PCB binding efficiency was calculated by using the equation: % PCB binding = [(A650/ε650)+(A539/ε539)]x(ε280/A280).

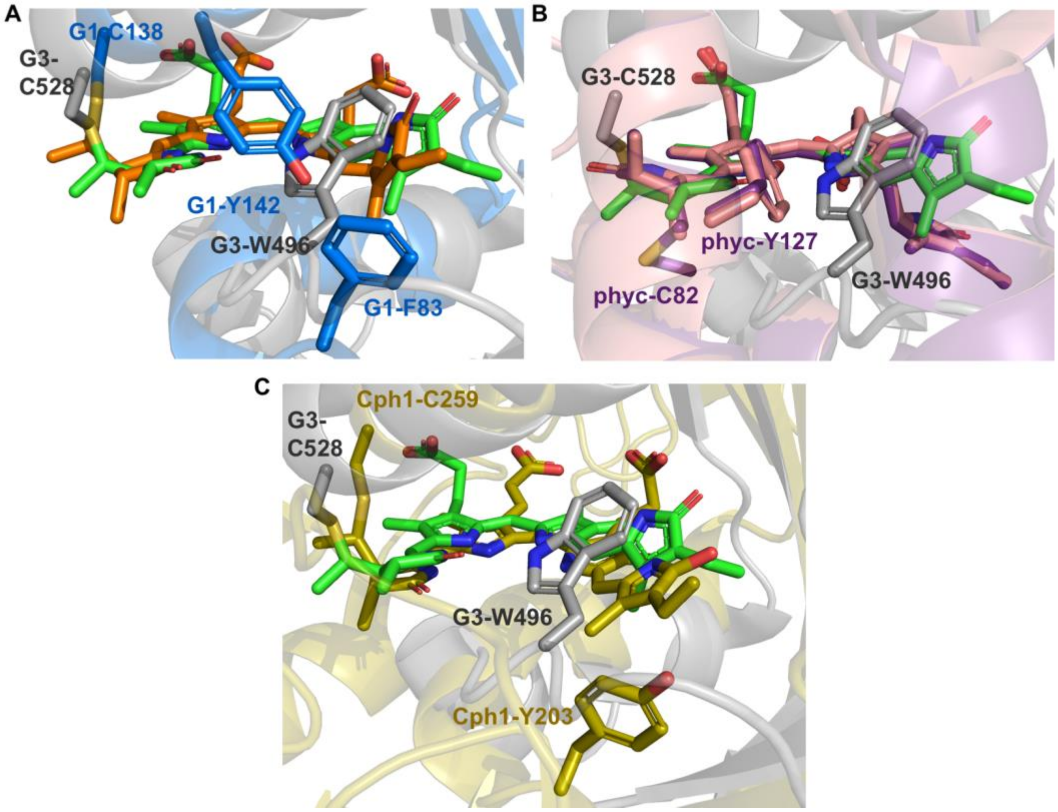


**Figure S3.** Crystal structure overlays of Slr1393 GAF3-PCB with PEB-bound proteins and Cph1-PCB. Overlays of PyMOL model of the crystal structure of Slr1393 GAF3 bound to PCB in the red state (PDB 5DFX) with A) PyMOL model of the crystal structure of All2699 GAF1 bound to PEB (PDB 6OZB), B) PyMOL models of the crystal structures of Slr1393 GAF3 bound to PCB in the red state (PDB 5DFX), B-phycoerythrin bound to PEB (PDB 3V58), and phycoerythrin bound to PEB (PDB 2VJH), and C) PyMOL model of the crystal structure of Cph1 bound to PCB (PDB 2VEA). The protein structure of GAF3 is colored gray, the PCB cofactor is green, and the side chain of the tryptophan 496 residue mutated in this work is shown as a gray stick. The protein structure of GAF1 is colored blue, the PEB cofactor orange, and the tyrosine 142 residue is shown as a blue stick. The phenylalanine 83 residue that aligns with W496 in the sequence alignment (Figure S4) is also shown as a blue stick. The protein structure of B-phycoerythrin (PDB3V58) is colored purple, the PEB cofactor is purple, and the side chain of the tyrosine 127 residue is shown as a purple stick. The protein structure of phycoerythrin (PDB 2VJH) is colored pink, the PEB cofactor is pink, and the side chain of the tyrosine 127 residue is shown as a pink stick. The protein structure of Cph1 is colored gold, the PCB cofactor gold, and the tyrosine 203 residue is shown as a gold stick.

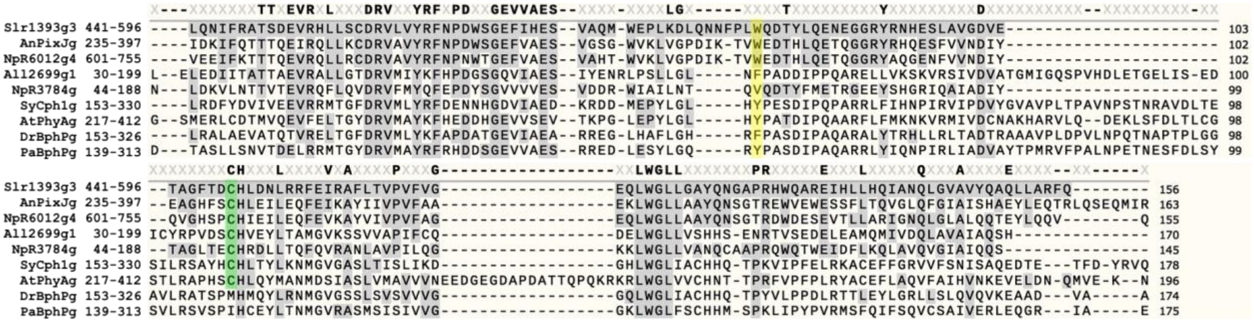


**Figure S4.** Multiple sequence alignment of GAF3 with other CBCRs and the GAF domains of phytochromes. The position of the W496 residue in GAF3 (Slr1393g3) is highlighted in yellow along with the corresponding residue in the other GAF domains. The position of the C528 residue in GAF3 that covalently binds at ring A of the bilin is highlighted in green and is conserved in most GAF domains except the phytochrome GAF domains DrBphPg and PaBphPg, where the Cys residue is located in a different domain (not shown). The UniProtKB IDs for each sequence are listed as follows: Slr1393g3: P73184; AnPixJg: Q8YXY7; NpR6012g4: B2IU14; All2699g1: Q8YTL8; NpR3784g: B2J457; SyCph1g: Q55168; AtPhyAg: P14712; DrBphPg: Q9RZA4; PaBphPg: Q9HWR3. This multiple sequence alignment was generated in SnapGene using T Coffee.

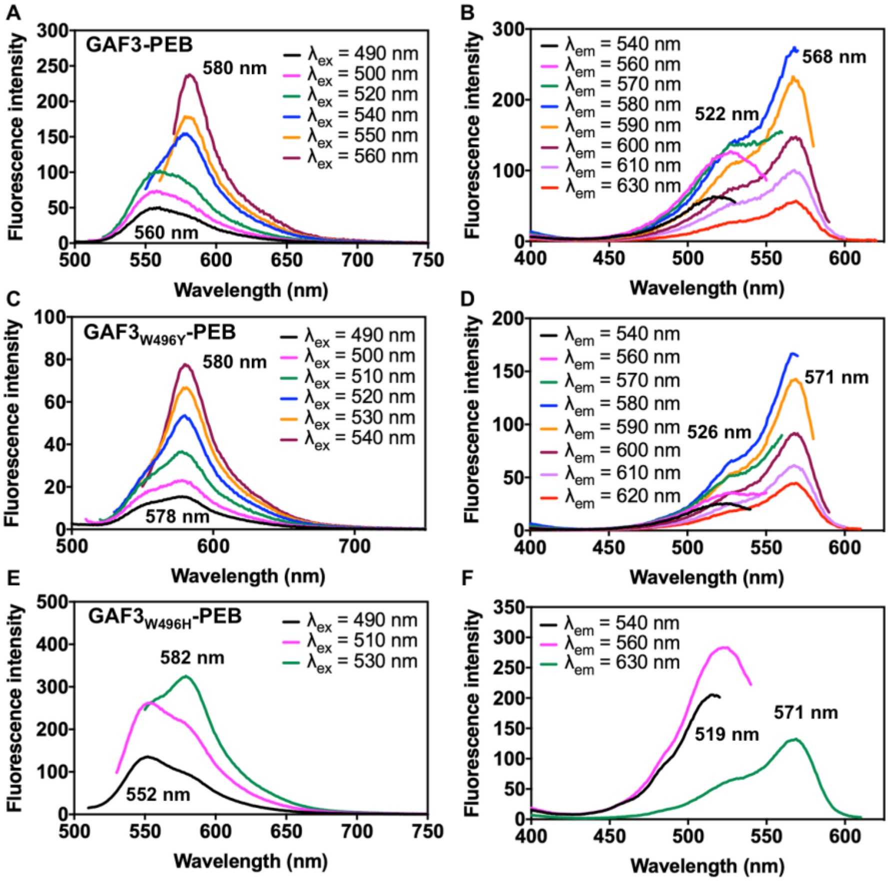


**Figure S5.** Fluorescence excitation and emission spectra of GAF3 and GAF3 mutants. Fluorescence emission spectra of GAF3-PEB (A), GAF3$_{W496Y}$-PEB (C), and GAF3$_{W496H}$-PEB (E) collected at various excitation wavelengths. Fluorescence excitation spectra of GAF3-PEB (B), GAF3$_{W496Y}$-PEB (D), and GAF3$_{W496H}$-PEB (F) collected at various emission wavelengths. Representative maximum wavelengths are indicated on each graph. Protein concentration = ~3 µM. Buffer is 50 mM HEPES, 100 mM NaCl, pH 7.1.

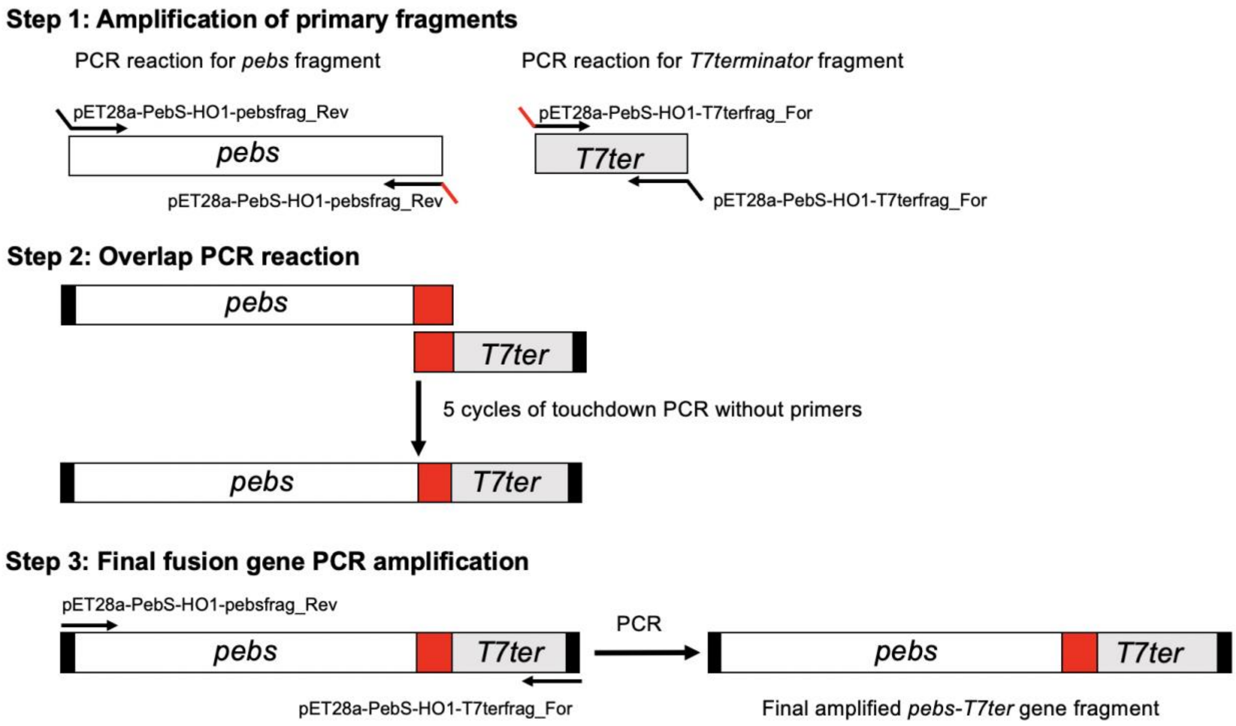


**Figure S6.** Schematic of PCR amplification to generate the pebs-T7ter fragment for plasmids pET28a-PebS-HO1 and pACYCDuet-PebS-T7-ter-HO1. Step 1 (top panel) uses touchdown PCR to generate PCR fragments that contain a 17 bp length overlap, which is represented by the red portion of primers pET28a-PebS-HO1-pebsfrag_Rev (Table S4) and pET28a-PebS-HO1-T7terfrag_For (Table S4), at the 3' end of the pebs fragment and the 5' end of the T7ter fragment. Step 2 (middle panel) ligates the pebs and T7ter fragments to generate a full-length pebs-T7ter gene fragment using overlap extension PCR (the annealing temperature is the Tm of the overlapping fragment, 60 °C). Step 3 demonstrates PCR amplification of the full-length pebs-T7ter fragment that was then used to generate plasmids pET28a-PebS-HO1 and pACYCDuet-PebS-T7-ter-HO1 as described in the Experimental Section above. All primers for generating this fragment are listed in Table S4.

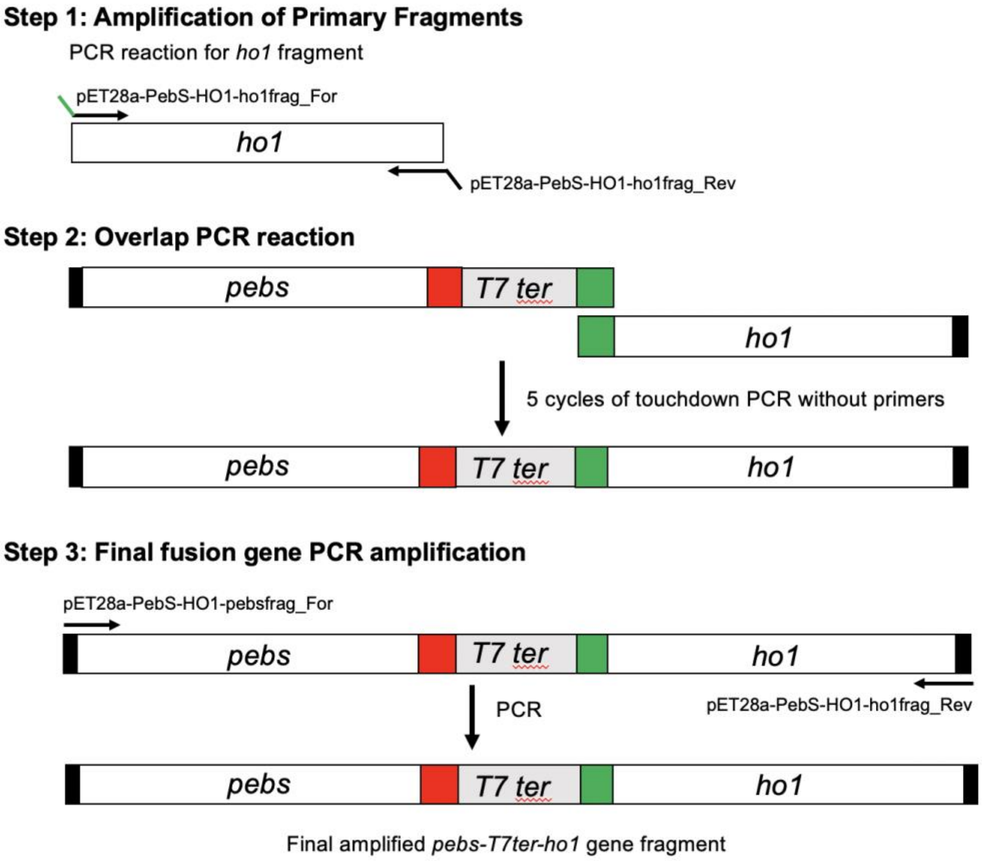


**Figure S7.** Schematic of PCR amplification to generate the *pebS-T7ter-ho1* fragment for plasmids pET28a-PebS-HO1 and pACYCDuet-PebS-T7-ter-HO1. Step 1 (top panel) uses touchdown PCR to generate a fragment that contains a 16 bp length overlap at the 5' end of the *ho1* gene fragment and the 3' end of the *pebs-T7ter* fragment (from Figure S4). Step 2 (middle panel) ligates the *ho1* and *pebs-T7ter* fragments to generate a full-length *pebs-T7ter-ho1* gene fragment using overlap extension PCR. The annealing temperature is the Tm of the overlapping fragment (58 °C, this overlap is represented by the green boxes). Step 3 demonstrates PCR amplification of the full length *pebS-T7ter-ho1* fragment that was then used to generate plasmids pET28a-PebS-HO1 and pACYCDuet-PebS-T7-ter-HO1 as described in the Experimental Section above. All primers for generating this fragment are listed in Table S4.

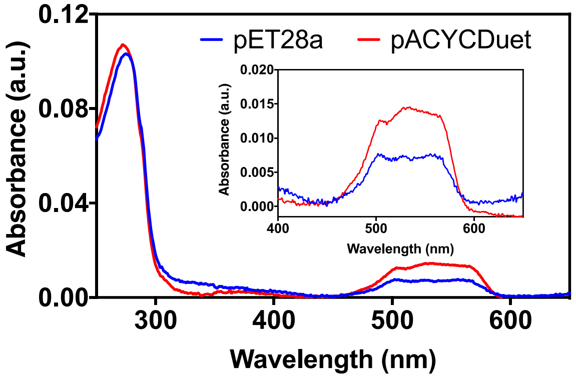


**Figure S8.** UV-visible absorption spectra of GAF3$_{W496Y}$-PEB produced when the protein was encoded in pET28a (high copy number) compared to pACYCDuet (low copy number). The region demonstrating bound PEB/PUB is enlarged in the inset. Protein concentration = 3 μM. Buffer = 50 mM HEPES, 100 mM NaCl, pH 7.1.

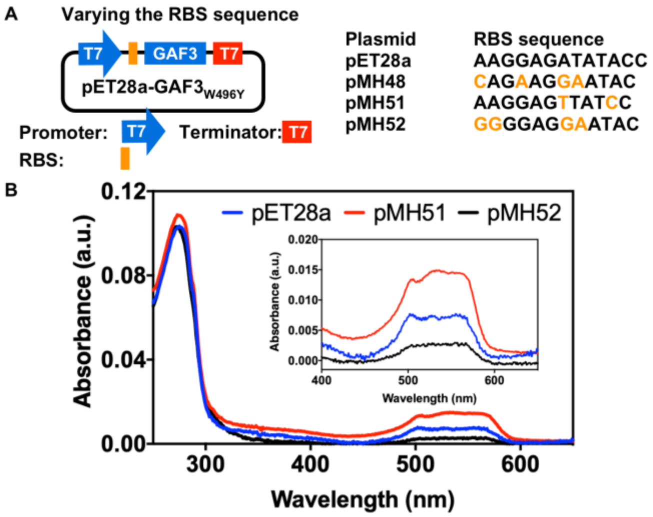


**Figure S9.** Plasmid designs and UV-visible spectra of $GAF3_{W496Y}$-PEB produced from pET28a and pMH plasmids with varied RBS sequences. A) Plasmid map overview and varied RBS sequences in pET28a, pMH48, pMH51, and pMH52. B) UV-visible absorption spectra of $GAF3_{W496Y}$-PEB produced when the protein was encoded in pET28a (original RBS sequence) compared to pMH plasmids (varied RBS sequences). The region demonstrating bound PEB/PUB is enlarged in the inset. Protein concentration = ~3 μM. Buffer is 50 mM HEPES, 100 mM NaCl, pH 7.1.

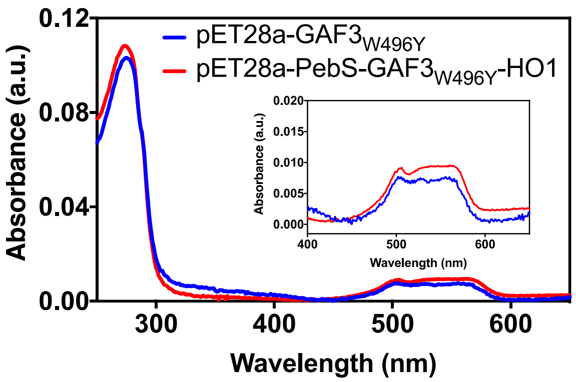


**Figure S10.** UV-visible absorption spectra of GAF3W496Y-PEB produced from pET28a-PebS-GAF3W496Y-HO1 compared to GAF3W496Y-PEB expressed from pET28a with coexpression of PEB synthesis enzymes. The region demonstrating bound PEB/PUB is enlarged in the inset. Protein concentration = ~3 μM. Buffer is 50 mM HEPES, 100 mM NaCl, pH 7.1.

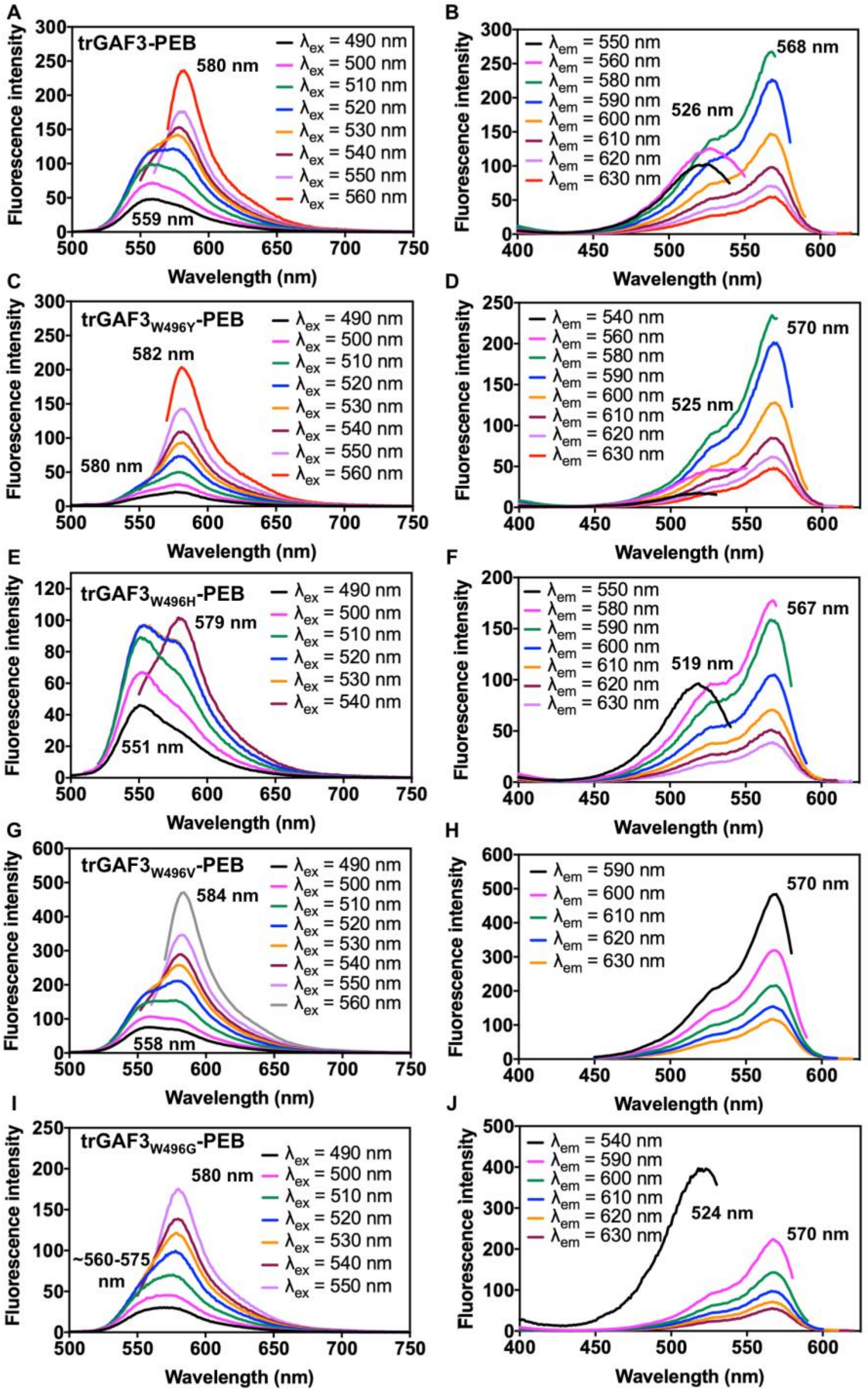


**Figure S11.** Fluorescence emission and excitation spectra for truncated GAF3 and mutants. Fluorescence emission spectra of trGAF3-PEB (A), trGAF3$_{W496Y}$-PEB (C), trGAF3$_{W496H}$-PEB (E), trGAF3$_{W496V}$-PEB (G), and trGAF3$_{W496G}$-PEB (I) collected at various excitation wavelengths. Fluorescence excitation spectra of trGAF3-PEB (B), trGAF3$_{W496Y}$-PEB (D), trGAF3$_{W496H}$-PEB (F), trGAF3$_{W496V}$-PEB (H), and trGAF3$_{W496G}$-PEB (J) collected at various emission wavelengths. Representative maximum wavelengths are indicated on each graph. Protein concentration = ~3 μM. Buffer is 50 mM HEPES, 100 mM NaCl, pH 7.1.

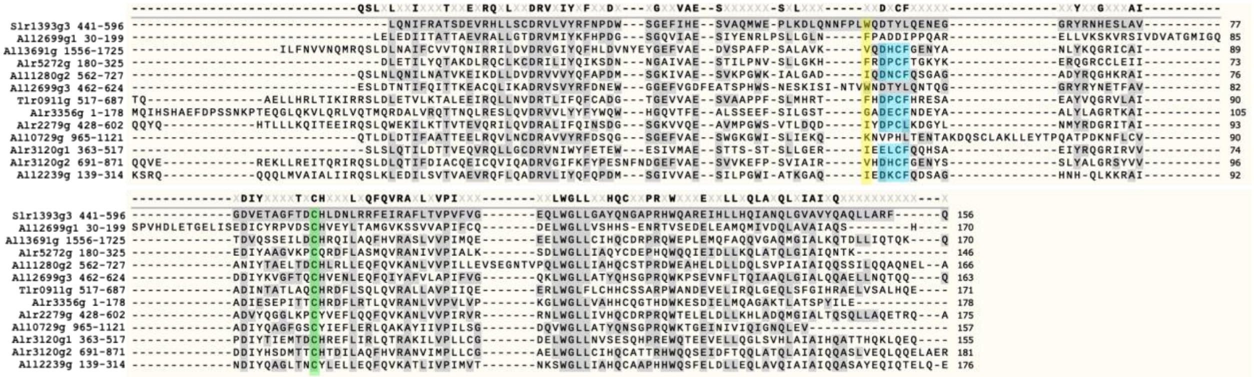


**Figure S12.** Multiple sequence alignment of GAF3 with other CBCRs tested for PEB binding.[4] The position of the W496 residue in GAF3 (Slr1393g3) is highlighted in yellow along with the corresponding residue in the other GAF domains. The position of the C528 residue in GAF3 that covalently binds at ring A of the bilin is highlighted in green and is conserved in these GAF domains. The DXCF motif (or similar) conserved among blue/green photoswitching CBCRs is highlighted in blue. In PCB-binding GAF domains containing this motif, the Cys residue reversibly binds to the C10 position of the cofactor, breaking the conjugation of the cofactor and allowing for blue/green photoswitching.[5,6] The domain All2699g1 binds PEB without autoisomerization. All3691g, Alr5272g, All1280g2, All2699g3, Tlr0911g, Alr3356g, and Alr2279g all bind PEB with partial autoisomerization to PUB. All0729g, Alr3120g1, Alr3120g2, and All2239g do not bind PEB. The UniProtKB IDs for each sequence are listed as follows: Slr1393g3: P73184; All2699g1 and All2699g3: Q8YTL8; All3691g: Q8YQX0; Alr5272g: Q8YLM6; All1280g2: Q8YXD3; Tlr0911g: Q8DKE8; Alr3356g: Q8YRT7; Alr2279g: Q8YUQ6; All0729g: Q8YYW3; Alr3120g1 and Alr3120g2: Q8YSG5; All2239g: Q8YUU2. This multiple sequence alignment was generated in SnapGene using T Coffee.

**pET28a-GAF3**

```
TAATACGACTCACTATAGGGGAATTGTGAGCGGATAACAATTCCCCTCTAGAAATAATTT
TGTTTAACTTTAAGAAGGAGATATACCATGGGCAGCAGCCATCATCATCATCATCACAGC
AGCGGCCTGGTGCCGCGCGGCAGCCATATGGCTAGCATGACTGGTGGACAGCAAATGGGT
CGCGGATCCTTACAGAATATTTTTCGGGCCACCAGTGATGAGGTGCGTCATTTGCTCAGT
TGTGACCGGGTGTTGGTCTATCGCTTTAATCCGGACTGGAGCGGGGAATTTATCCATGAG
TCGGTGGCCCAAATGTGGGAACCACTGAAAGACTTGCAAAATAATTTCCCCCTCTGGCAA
GACACCTATCTCCAGGAAAATGAGGGGGGTCGTTACCGCAACCATGAGAGTTTGGCGGTG
GGGGACGTGGAAACAGCCGGTTTTACCGATTGCCATTTGGACAATTTGCGCCGGTTTGAA
ATTCGTGCCTTTTTAACTGTGCCAGTGTTTGTGGGGGAACAACTCTGGGGCCTGTTGGGG
GCCTATCAAAATGGGGCTCCCCGCCATTGGCAAGCGAGGGAAATCCATTTGCTCCATCAA
ATTGCCAATCAGTTGGGCGTAGCAGTGTACCAAGCCCAGTTGCTGGCCCGCTTCCAATGA
CTCGAGCACCACCACCACCACCACTGAGATCCGGCTGCTAACAAAGCCCGAAAGGAAGCT
GAGTTGGCTGCTGCCACCGCTGAGCAATAACTAGCATAACCCCTTGGGGCCTCTAAACGG
GTCTTGAGGGGTTTTTTG
```

T7 promoter

RBS 6xHis tag

Thrombin site T7 tag

BamHI GAF3 Trp496

XhoI

T7 terminator

**Figure S13.** Nucleotide sequence of the promoter and coding regions of pET28a-GAF3. The T7 promoter, RBS sequence, T7 terminator, and coding regions for a $His_6$ tag, thrombin cleavage site, T7 tag, and GAF3 are all highlighted as shown. The restriction sites used for cloning GAF3 into the pET28a backbone are also highlighted as shown. The coding sequence for the Trp residue in position 496 of the GAF3 protein is colored yellow.

**pET28a-GAF3$_{W496Y}$**

```
TAATACGACTCACTATAGGGGAATTGTGAGCGGATAACAATTCCCCTCTAGAAATAATTT
TGTTTAACTTTAAGAAGGAGATATACCATGGGCAGCAGCCATCATCATCATCATCACAGC
AGCGGCCTGGTGCCGCGCGGCAGCCATATGGCTAGCATGACTGGTGGACAGCAAATGGGT
CGCGGATCCTTACAGAATATTTTTCGGGCCACCAGTGATGAGGTGCGTCATTTGCTCAGT
TGTGACCGGGTGTTGGTCTATCGCTTTAATCCGGACTGGAGCGGGGAATTTATCCATGAG
TCGGTGGCCCAAATGTGGGAACCACTGAAAGACTTGCAAAATAATTTCCCCCTCTATCAA
GACACCTATCTCCAGGAAAATGAGGGGGGTCGTTACCGCAACCATGAGAGTTTGGCGGTG
GGGGACGTGGAAACAGCCGGTTTTACCGATTGCCATTTGGACAATTTGCGCCGGTTTGAA
ATTCGTGCCTTTTTAACTGTGCCAGTGTTTGTGGGGGAACAACTCTGGGGCCTGTTGGGG
GCCTATCAAAATGGGGCTCCCCGCCATTGGCAAGCGAGGGAAATCCATTTGCTCCATCAA
ATTGCCAATCAGTTGGGCGTAGCAGTGTACCAAGCCCAGTTGCTGGCCCGCTTCCAATGA
CTCGAGCACCACCACCACCACCACTGAGATCCGGCTGCTAACAAAGCCCGAAAGGAAGCT
GAGTTGGCTGCTGCCACCGCTGAGCAATAACTAGCATAACCCCTTGGGGCCTCTAAACGG
GTCTTGAGGGGTTTTTTG
```

T7 promoter
RBS 6xHis tag
Thrombin site T7 tag
BamHI GAF3 Tyr496
XhoI
T7 terminator

**Figure S14.** Nucleotide sequence of the promoter and coding regions of pET28a-GAF3$_{W496Y}$. This plasmid was prepared by site-directed mutagenesis of pET28a-GAF3 (Figure S7, Table S2) at position 496. The T7 promoter, RBS sequence, T7 terminator, and coding regions for a $His_6$ tag, thrombin cleavage site, T7 tag, and GAF3 are all highlighted as shown. The restriction sites used for cloning GAF3 into the pET28a backbone are also highlighted as shown. The coding sequence for the mutated Trp to Tyr residue in position 496 of the GAF3 protein is colored yellow.

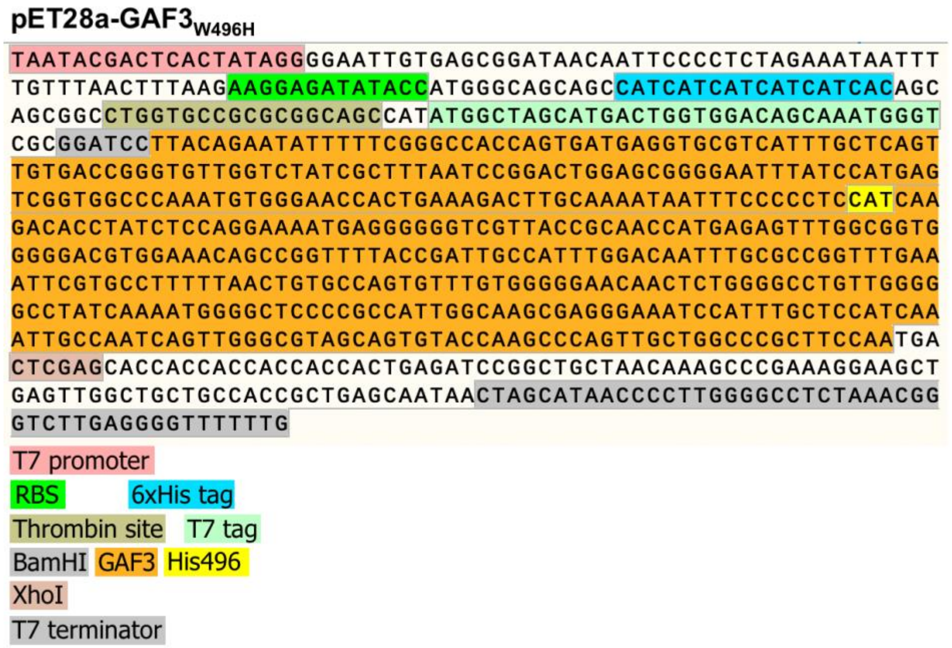


**Figure S15.** Nucleotide sequence of the promoter and coding regions of pET28a-GAF3$_{W496H}$. This plasmid was prepared by site-directed mutagenesis of pET28a-GAF3 (Figure S7, Table S2) at position 496. The T7 promoter, RBS sequence, T7 terminator, and coding regions for a His$_6$ tag, thrombin cleavage site, T7 tag, and GAF3 are all highlighted as shown. The restriction sites used for cloning GAF3 into the pET28a backbone are also highlighted as shown. The coding sequence for the mutated Trp to His residue in position 496 of the GAF3 protein is colored yellow.

**pACYCDuet-HO1-PcyA**

```
TAATACGACTCACTATAGGGGAATTGTGAGCGGATAACAATTCCCCTGTAGAAATAATTT
TGTTTAACTTTAATAAGGAGATATACCATGGGCAGCAGCCATCACCATCATCACCACAGC
CAGGATCCGAATTCCatgagtgtcaacttagcttcccagttgcgggaagggacgaaaaaa
tcccactccatggcggagaacgtcggctttgtcaaatgcttcctcaagggcgttgtcgag
aaaaattcctaccgtaagctggttggcaatctctactttgtctacagtgccatggaagag
gaaatggcaaaatttaaggaccatcccatcctcagccacatttacttccccgaactcaac
cgcaaacaaagcctagagcaagacctgcaattctattacggctccaactggcggcaagaa
gtgaaaatttctgccgctggccaagcctatgtggaccgagtccggcaagtggccgctacg
gcccctgaattgttggtggcccattcctacacccgttacctggggggatctttccggcggt
caaattctcaagaaaattgcccaaaatgccatgaatctccacgatggtggcacagctttc
tatgaatttgccgacattgatgacgaaaaggcttttaaaaatacctaccgtcaagctatg
aatgatctgcccattgaccaagccaccgccgaacggattgtggatgaagccaatgacgcc
tttgccatgaacatgaaaatgttcaacgaacttgaaggcaacctgatcaaggcgatcggc
attatggtgttcaacagcctcacccgtcgccgcagtcaaggcagcaccgaagttggcctc
gccacctccgaaggctagttaaagaggagaaaggatccatggccgtcactgatttaagtt
tgaccaattcttccctgatgcccacgttgaacccgatgattcaacagttggccctggcga
tcgccgctagttggcaaagtttacccctcaagccctatcaattgccggaggatttgggct
acgtagaaggccgcctggaaggggaaaagttagtgattgaaaatcggtgctaccaaacgc
cccagtttcgcaaaatgcatttggagttggccaaggtgggcaaagggttggatattctcc
actgtgtaatgtttcctgagcctttatacggtctacctttgtttggctgtgacattgtgg
ccggccccggtggagtaagtgcggctattgcggatctatcccccacccaaagcgatcgcc
aattgcccgcagcgtaccaaaaatcattggcagagctaggccagccagaatttgagcaac
aacgggaattgccccccctggggagaaatatttttctgaatattgtttattcatccgtccca
gcaatgtcactgaagaagaaagatttgtacaaagggtagtggacttttttgcaaattcatt
gtcaccaatccatcgttgccgaacccttgtctgaagctcaaactttggagcaccgtcagg
ggcaaattcattactgccaacaacaacagaaaaatgataaaacccgtcgggtactggaaa
aagcttttggggaagcttgggcggaacggtatatgagccaagtcttatttgatgttatcc
aataaGAGCTC
```

T7 promoter

RBS 6xHis tag

EcoRI HO1 PcyA

SacI

**Figure S16.** Nucleotide sequence of the promoter and coding regions of pACYCDuet-HO1-PcyA. The T7 promoter, RBS sequences, and coding regions for a $His_6$ tag, HO1, and PcyA are all highlighted as shown. The restriction sites used for cloning HO1-PcyA into the pACYCDuet-1 backbone are also highlighted as shown.

**pACYCDuet-PebS-HO1**

```
TAATACGACTCACTATAGGGGAATTGTGAGCGGATAACAATTCCCCTGTAGAAATAATTT
TGTTTAACTTTAATAAGGAGATATACCATGGcaggatccATGACAAAGAACCCACGCAAT
AATAAACCCAAGAAGATCTTGGACAGCAGCTATAAGTCCAAGACGATCTGGCAGAACTAT
ATCGACGCTCTGTTCGAAACCTTCCCGCAGTTGGAAATTAGCGAAGTGTGGGCGAAATGG
GATGGTGGTAATGTCACCAAAGATGGCGGCGATGCCAAGCTGACCGCAAATATCCGTACC
GGTGAGCACTTCTTAAAGGCGCGTGAAGCGCATATTGTGGACCCTAATAGCGACATCTAC
AACACCATCCTGTATCCGAAAACCGGTGCAGATCTTCCGTGTTTTGGCATGGATCTGATG
AAATTTTCCGACAAAAAAGTTATTATTGTTTTTGATTTCCAACATCCGCGCGAGAAATAC
TTGTTCTCTGTTGACGGCCTGCCGGAAGATGATGGCAAATACCGCTTTTTTGAGATGGGT
AACCACTTTTCGAAAAACATCTTTGTGCGCTACTGCAAACCGGATGAAGTCGATCAATAT
CTCGACACCTTCAAACTGTACCTGACGAAGTACAAGGAGATGATTGACAACAACAAGCCA
GTGGGTGAGGACACGACCGTTTATAGCGACTTCGACACTTACATGACCGAACTGGACCCG
GTGCGTGGTTATATGAAGAACAAGTTCGGCGAGGGTCGTAGCGAGGCGTTTGTTAATGAT
TTCCTGTTCTCTTACAAGTAAGAGCTCGGCGCGCCTGCAGGTCGACAAGCTTGCGGCCGC
ATAATGCTTAAGTCGAACAGAAAGTAATCGTATTGTACACGGCCGCATAATCGAAATTAA
TACGACTCACTATAGGGGAATTGTGAGCGGATAACAATTCCCCATCTTAGTATATTAGTT
AAGTATAAGAAGGAGATATACATATGggcatgagtgtcaacttagcttcccagttgcggg
aagggacgaaaaaatcccactccatggcggagaacgtcggctttgtcaaatgcttcctca
agggcgttgtcgagaaaaattcctaccgtaagctggttggcaatctctactttgtctaca
gtgccatggaagaggaaatggcaaaatttaaggaccatcccatcctcagccacatttact
tccccgaactcaaccgcaaacaaagcctagagcaagacctgcaattctattacggctcca
actggcggcaagaagtgaaaatttctgccgctggccaagcctatgtggaccgagtccggc
aagtggccgctacggcccctgaattgttggtggcccattcctacacccgttacctgggg
atctttccggcggtcaaattctcaagaaaattgcccaaaatgccatgaatctccacgatg
gtggcacagctttctatgaatttgccgacattgatgacgaaaaggcttttaaaaatacct
accgtcaagctatgaatgatctgcccattgaccaagccaccgccgaacggattgtggatg
aagccaatgacgcctttgccatgaacatgaaaatgttcaacgaacttgaaggcaacctga
tcaaggcgatcggcattatggtgttcaacagcctcacccgtcgccgcagtcaaggcagca
ccgaagttggcctcgccacctccgaaggctagCTCGAGTCTGGTAAAGAAACCGCTGCTG
CGAAATTTGAACGCCAGCACATGGACTCGTCTACTAGCGCAGCTTAATTAACCTAGGCTG
CTGCCACCGCTGAGCAATAACTAGCATAACCCCTTGGGGCCTCTAAACGGGTCTTGAGGG
GTTTTTTG
```

T7 promoter

RBS    PebS

NdeI    HO1

XhoI

T7 terminator

**Figure S17.** Nucleotide sequence of the promoter and coding regions of pACYCDuet-PebS-HO1. The T7 promoter, RBS sequences, T7 terminator, and coding regions for PebS and HO1 are all highlighted as shown. The restriction sites used for cloning HO1 into the pACYCDuet-1 backbone (prior to NEB HiFi DNA assembly for insertion of PebS) are also highlighted as shown.

**pET28a-PebS-HO1**

```
TAATACGACTCACTATAGGGGAATTGTGAGCGGATAACAATTCCCCTCTAGAAATAATTT
TGTTTAACTTTAAGAAGGAGATATACCATGACAAAGAACCCACGCAATAATAAACCCAAG
AAGATCTTGGACAGCAGCTATAAGTCCAAGACGATCTGGCAGAACTATATCGACGCTCTG
TTCGAAACCTTCCCGCAGTTGGAAATTAGCGAAGTGTGGGCGAAATGGGATGGTGGTAAT
GTCACCAAAGATGGCGGCGATGCCAAGCTGACCGCAAATATCCGTACCGGTGAGCACTTC
TTAAAGGCGCGTGAAGCGCATATTGTGGACCCTAATAGCGACATCTACAACACCATCCTG
TATCCGAAAACCGGTGCAGATCTTCCGTGTTTTGGCATGGATCTGATGAAATTTTCCGAC
AAAAAAGTTATTATTGTTTTTGATTTCCAACATCCGCGCGAGAAATACTTGTTCTCTGTT
GACGGCCTGCCGGAAGATGATGGCAAATACCGCTTTTTTGAGATGGGTAACCACTTTTCG
AAAAACATCTTTGTGCGCTACTGCAAACCGGATGAAGTCGATCAATATCTCGACACCTTC
AAACTGTACCTGACGAAGTACAAGGAGATGATTGACAACAACAAGCCAGTGGGTGAGGAC
ACGACCGTTTATAGCGACTTCGACACTTACATGACCGAACTGGACCCGGTGCGTGGTTAT
ATGAAGAACAAGTTCGGCGAGGGTCGTAGCGAGGCGTTTGTTAATGATTTCCTGTTCTCT
TACAAGTAAGATCCGGCTGCTAACAAAGCCCGAAAGGAAGCTGAGTTGGCTGCTGCCACC
GCTGAGCAATAACTAGCATAACCCCTTGGGGCCTCTAAACGGGTCTTGAGGGGTTTTTTG
GAGCTCGGCGCGCCTGCAGGTCGACAAGCTTGCGGCCGCATAATGCTTAAGTCGAACAGA
AAGTAATCGTATTGTACACGGCCGCATAATCGAAATTAATACGACTCACTATAGGGGAAT
TGTGAGCGGATAACAATTCCCCATCTTAGTATATTAGTTAAGTATAAGAAGGAGATATAC
ATATGggcatgagtgtcaacttagcttcccagttgcgggaagggacgaaaaaatcccact
ccatggcggagaacgtcggctttgtcaaatgcttcctcaagggcgttgtcgagaaaaatt
cctaccgtaagctggttggcaatctctactttgtctacagtgccatggaagaggaaatgg
caaaatttaaggaccatcccatcctcagccacatttacttccccgaactcaaccgcaaac
aaagcctagagcaagacctgcaattctattacggctccaactggcggcaagaagtgaaaa
tttctgccgctggccaagcctatgtggaccgagtccggcaagtggccgctacggcccctg
aattgttggtggcccattcctacacccgttacctggggatctttccggcggtcaaattc
tcaagaaaattgcccaaaatgccatgaatctccacgatggtggcacagctttctatgaat
ttgccgacattgatgacgaaaaggcttttaaaaatacctaccgtcaagctatgaatgatc
tgcccattgaccaagccaccgccgaacggattgtggatgaagccaatgacgcctttgcca
tgaacatgaaaatgttcaacgaacttgaaggcaacctgatcaaggcgatcggcattatgg
tgttcaacagcctcacccgtcgccgcagtcaaggcagcaccgaagttggcctcgccacct
ccgaaggctaaGATCCGGCTGCTAACAAAGCCCGAAAGGAAGCTGAGTTGGCTGCTGCCA
CCGCTGAGCAATAACTAGCATAACCCCTTGGGGCCTCTAAACGGGTCTTGAGGGGTTTTT
TG
```

T7 promoter
RBS PebS
T7 terminator HO1

**Figure S18.** Nucleotide sequence of the promoter and coding regions of pET28a-PebS-HO1. The T7 promoters, RBS sequences, T7 terminators, and coding regions for PebS and HO1 are all highlighted as shown.

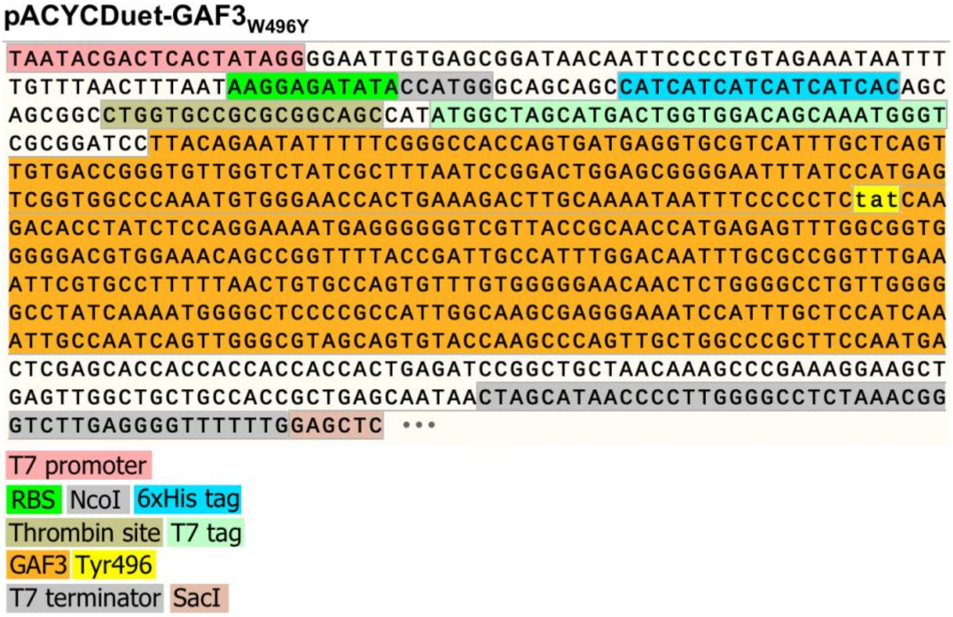


**Figure S19.** Nucleotide sequence of the promoter and coding regions of pACYCDuet-GAF3$_{W496Y}$. The T7 promoter, RBS sequence, T7 terminator, and coding regions for a His$_6$ tag, thrombin cleavage site, T7 tag, and GAF3 are all highlighted as shown. The restriction sites used for cloning GAF3 into the pACYCDuet backbone are also highlighted as shown. The coding sequence for the mutated Trp to His residue in position 496 of the GAF3 protein is colored yellow.

**pACYCDuet-PebS-T7-ter-HO1**

```
TAATACGACTCACTATAGGGGAATTGTGAGCGGATAACAATTCCCCTGTAGAAATAATTT
TGTTTAACTTTAATAAGGAGATATACCATGACAAAGAACCCACGCAATAATAAACCCAAG
AAGATCTTGGACAGCAGCTATAAGTCCAAGACGATCTGGCAGAACTATATCGACGCTCTG
TTCGAAACCTTCCCGCAGTTGGAAATTAGCGAAGTGTGGGCGAAATGGGATGGTGGTAAT
GTCACCAAAGATGGCGGCGATGCCAAGCTGACCGCAAATATCCGTACCGGTGAGCACTTC
TTAAAGGCGCGTGAAGCGCATATTGTGGACCCTAATAGCGACATCTACAACACCATCCTG
TATCCGAAAACCGGTGCAGATCTTCCGTGTTTTGGCATGGATCTGATGAAATTTTCCGAC
AAAAAAGTTATTATTGTTTTTGATTTCCAACATCCGCGCGAGAAATACTTGTTCTCTGTT
GACGGCCTGCCGGAAGATGATGGCAAATACCGCTTTTTTGAGATGGGTAACCACTTTTCG
AAAAACATCTTTGTGCGCTACTGCAAACCGGATGAAGTCGATCAATATCTCGACACCTTC
AAACTGTACCTGACGAAGTACAAGGAGATGATTGACAACAACAAGCCAGTGGGTGAGGAC
ACGACCGTTTATAGCGACTTCGACACTTACATGACCGAACTGGACCCGGTGCGTGGTTAT
ATGAAGAACAAGTTCGGCGAGGGTCGTAGCGAGGCGTTTGTTAATGATTTCCTGTTCTCT
TACAAGTAAGATCCGGCTGCTAACAAAGCCCGAAAGGAAGCTGAGTTGGCTGCTGCCACC
GCTGAGCAATAACTAGCATAACCCCTTGGGGCCTCTAAACGGGTCTTGAGGGGTTTTTTG
GAGCTCGGCGCGCCTGCAGGTCGACAAGCTTGCGGCCGCATAATGCTTAAGTCGAACAGA
AAGTAATCGTATTGTACACGGCCGCATAATCGAAATTAATACGACTCACTATAGGGGAAT
TGTGAGCGGATAACAATTCCCCATCTTAGTATATTAGTTAAGTATAAGAAGGAGATATAC
ATATGggcatgagtgtcaacttagcttcccagttgcgggaagggacgaaaaaatcccact
ccatggcggagaacgtcggctttgtcaaatgcttcctcaagggcgttgtcgagaaaaatt
cctaccgtaagctggttggcaatctctactttgtctacagtgccatggaagaggaaatgg
caaaatttaaggaccatcccatcctcagccacatttacttccccgaactcaaccgcaaac
aaagcctagagcaagacctgcaattctattacggctccaactggcggcaagaagtgaaaa
tttctgccgctggccaagcctatgtggaccgagtccggcaagtggccgctacggcccctg
aattgttggtggcccattcctacacccgttacctggggggatctttccggcggtcaaattc
tcaagaaaattgcccaaaatgccatgaatctccacgatggtggcacagctttctatgaat
ttgccgacattgatgacgaaaaggcttttaaaaatacctaccgtcaagctatgaatgatc
tgcccattgaccaagccaccgccgaacggattgtggatgaagccaatgacgcctttgcca
tgaacatgaaaatgttcaacgaacttgaaggcaacctgatcaaggcgatcggcattatgg
tgttcaacagcctcacccgtcgccgcagtcaaggcagcaccgaagttggcctcgccacct
ccgaaggctaaCTCGAGTCTGGTAAAGAAACCGCTGCTGCGAAATTTGAACGCCAGCACA
TGGACTCGTCTACTAGCGCAGCTTAATTAACCTAGGCTGCTGCCACCGCTGAGCAATAAC
TAGCATAACCCCTTGGGGCCTCTAAACGGGTCTTGAGGGGTTTTTTG
```

T7 promoter
RBS PebS
T7 terminator HO1

**Figure S20.** Nucleotide sequence of the promoter and coding regions of pACYCDuet-PebS-T7-ter-HO1. The T7 promoters, RBS sequences, T7 terminators, and coding regions for PebS and HO1 are all highlighted as shown.

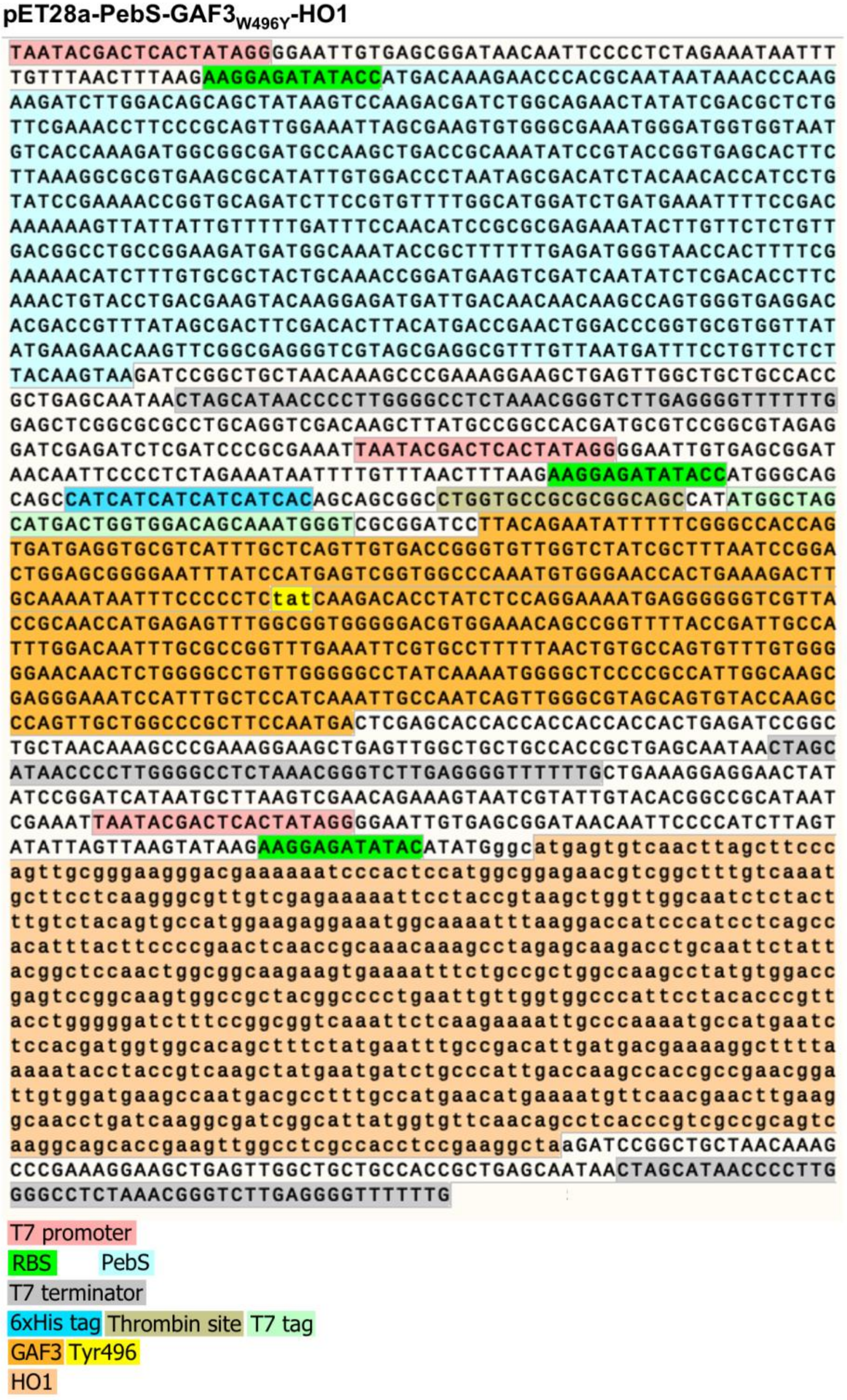


**Figure S21.** Nucleotide sequence of the promoter and coding regions of pET28a-PebS-GAF3$_{W496Y}$-HO1. The T7 promoters, RBS sequences, T7 terminators and coding regions for a His$_6$ tag, thrombin cleavage site, T7 tag, PebS, GAF3$_{W496Y}$, and HO1 are all highlighted as shown. The coding sequence for the mutated Trp to Tyr residue in position 496 of the GAF3 protein is colored yellow.

**pET28a-PebS-GAF3$_{W496Y}$-HO1**

```
TAATACGACTCACTATAGGGGAATTGTGAGCGGATAACAATTCCCCTCTAGAAATAATTT
TGTTTAACTTTAAGAAGGAGATATACCATGACAAAGAACCCACGCAATAATAAACCCAAG
AAGATCTTGGACAGCAGCTATAAGTCCAAGACGATCTGGCAGAACTATATCGACGCTCTG
TTCGAAACCTTCCCGCAGTTGGAAATTAGCGAAGTGTGGGCGAAATGGGATGGTGGTAAT
GTCACCAAAGATGGCGGCGATGCCAAGCTGACCGCAAATATCCGTACCGGTGAGCACTTC
TTAAAGGCGCGTGAAGCGCATATTGTGGACCCTAATAGCGACATCTACAACACCATCCTG
TATCCGAAAACCGGTGCAGATCTTCCGTGTTTTGGCATGGATCTGATGAAATTTTCCGAC
AAAAAAGTTATTATTGTTTTTGATTTCCAACATCCGCGCGAGAAATACTTGTTCTCTGTT
GACGGCCTGCCGGAAGATGATGGCAAATACCGCTTTTTTGAGATGGGTAACCACTTTTCG
AAAAACATCTTTGTGCGCTACTGCAAACCGGATGAAGTCGATCAATATCTCGACACCTTC
AAACTGTACCTGACGAAGTACAAGGAGATGATTGACAACAACAAGCCAGTGGGTGAGGAC
ACGACCGTTTATAGCGACTTCGACACTTACATGACCGAACTGGACCCGGTGCGTGGTTAT
ATGAAGAACAAGTTCGGCGAGGGTCGTAGCGAGGCGTTTGTTAATGATTTCCTGTTCTCT
TACAAGTAAGATCCGGCTGCTAACAAAGCCCGAAAGGAAGCTGAGTTGGCTGCTGCCACC
GCTGAGCAATAACTAGCATAACCCCTTGGGGCCTCTAAACGGGTCTTGAGGGGTTTTTTG
GAGCTCGGCGCGCCTGCAGGTCGACAAGCTTATGCCGGCCACGATGCGTCCGGCGTAGAG
GATCGAGATCTCGATCCCGCGAAATTAATACGACTCACTATAGGGGAATTGTGAGCGGAT
AACAATTCCCCTCTAGAAATAATTTTGTTTAACTTTAAGAAGGAGATATACCATGGGCAG
CAGCCATCATCATCATCATCACAGCAGCGGCCTGGTGCCGCGCGGCAGCCATATGGCTAG
CATGACTGGTGGACAGCAAATGGGTCGCGGATCCTTACAGAATATTTTTCGGGCCACCAG
TGATGAGGTGCGTCATTTGCTCAGTTGTGACCGGGTGTTGGTCTATCGCTTTAATCCGGA
CTGGAGCGGGGAATTTATCCATGAGTCGGTGGCCCAAATGTGGGAACCACTGAAAGACTT
GCAAAATAATTTCCCCCTCTATCAAGACACCTATCTCCAGGAAAATGAGGGGGGTCGTTA
CCGCAACCATGAGAGTTTGGCGGTGGGGGACGTGGAAACAGCCGGTTTTACCGATTGCCA
TTTGGACAATTTGCGCCGGTTTGAAATTCGTGCCTTTTTAACTGTGCCAGTGTTTGTGGG
GGAACAACTCTGGGGCCTGTTGGGGGCCTATCAAAATGGGGCTCCCCGCCATTGGCAAGC
GAGGGAAATCCATTTGCTCCATCAAATTGCCAATCAGTTGGGCGTAGCAGTGTACCAAag
cTGACTCGAGCACCACCACCACCACCACTGAGATCCGGCTGCTAACAAAGCCCGAAAGGA
AGCTGAGTTGGCTGCTGCCACCGCTGAGCAATAACTAGCATAACCCCTTGGGGCCTCTAA
ACGGGTCTTGAGGGGTTTTTTGCTGAAAGGAGGAACTATATCCGGATCATAATGCTTAAG
TCGAACAGAAAGTAATCGTATTGTACACGGCCGCATAATCGAAATTAATACGACTCACTA
TAGGGGAATTGTGAGCGGATAACAATTCCCCATCTTAGTATATTAGTTAAGTATAAGAAG
GAGATATACATATGggcatgagtgtcaacttagcttcccagttgcgggaagggacgaaaa
aatcccactccatggcggagaacgtcggctttgtcaaatgcttcctcaagggcgttgtcg
agaaaaattcctaccgtaagctggttggcaatctctactttgtctacagtgccatggaag
aggaaatggcaaaatttaaggaccatcccatcctcagccacatttacttccccgaactca
accgcaaacaaagcctagagcaagacctgcaattctattacggctccaactggcggcaag
aagtgaaaatttctgccgctggccaagcctatgtggaccgagtccggcaagtggccgcta
cggcccctgaattgttggtggcccattcctacacccgttacctgggggatctttccggcg
gtcaaattctcaagaaaattgcccaaaatgccatgaatctccacgatggtggcacagctt
tctatgaatttgccgacattgatgacgaaaaggcttttaaaaatacctaccgtcaagcta
tgaatgatctgcccattgaccaagccaccgccgaacggattgtggatgaagccaatgacg
cctttgccatgaacatgaaaatgttcaacgaacttgaaggcaacctgatcaaggcgatcg
gcattatggtgttcaacagcctcacccgtcgccgcagtcaaggcagcaccgaagttggcc
tcgccacctccgaaggctaaGATCCGGCTGCTAACAAAGCCCGAAAGGAAGCTGAGTTGG
CTGCTGCCACCGCTGAGCAATAACTAGCATAACCCCTTGGGGCCTCTAAACGGGTCTTGA
GGGGTTTTTTG
```

T7 promoter

RBS PebS

T7 terminator

6xHis tag Thrombin site T7 tag

trGAF3 Tyr496

HO1

**Figure S22.** Nucleotide sequence of the promoter and coding regions of pET28a-trGAF3$_{W496Y}$. The T7 promoters, RBS sequences, T7 terminators and coding regions for a His$_6$ tag, thrombin cleavage site, T7 tag, PebS, truncated GAF3$_{W496Y}$, and HO1 are all highlighted as shown. The coding sequence for the mutated Trp to Tyr residue in position 496 of the GAF3 protein is colored yellow.

### References


[1] S. R. Schmidl, R. U. Sheth, A. Wu, J. J. Tabor, *ACS Synth. Biol.* **2014**, *3*, 820–831.

[2] J. Zhang, X. J. Wu, Z. B. Wang, Y. Chen, X. Wang, M. Zhou, H. Scheer, K. H. Zhao, *Angew. Chem. Int. Ed.* **2010**, *49*, 5456–5458.

[3] J. Stiefelmaier, B. Ledermann, M. Sorg, A. Banek, D. Geib, R. Ulber, N. Frankenberg-Dinkel, *J. Biotechnol.* **2018**, *274*, 47–53.

[4] Y. F. Sun, J. G. Xu, K. Tang, D. Miao, W. Gärtner, H. Scheer, K. H. Zhao, M. Zhou, *Photochem. Photobiol. Sci.* **2014**, *13*, 757–763.

[5] N. C. Rockwell, S. S. Martin, J. C. Lagarias, *Biochemistry* **2012**, *51*, 3576–3585.

[6] N. C. Rockwell, S. S. Martin, A. G. Gulevich, J. C. Lagarias, *Biochemistry* **2012**, *51*, 1449–1463.
